# Transgenic tools targeting striatal and pallidal subpopulations revealed evolutionary conservation and specialization of the cortico-basal ganglia circuit in zebrafish

**DOI:** 10.1101/2023.09.20.558712

**Authors:** Yuki Tanimoto, Hisaya Kakinuma, Ryo Aoki, Toshiyuki Shiraki, Shin-ichi Higashijima, Hitoshi Okamoto

## Abstract

The cortico-basal ganglia circuit mediates decision-making. Here, we generated transgenic tools for adult zebrafish targeting specific subpopulations of the components of this circuit and utilized them to identify evolutionary homologs of the mammalian direct- and indirect-pathway striatal neurons which respectively project to the homologs of the internal and external segment of the globus pallidus (dEN and Vl) as in mammals. Unlike in mammals, the Vl mainly projected to the dEN directly, not by way of the subthalamic nucleus. Further single-cell RNA sequencing analysis revealed two pallidal output pathways: a major shortcut pathway directly connecting the dEN with the pallium and the evolutionarily conserved closed loop by way of the thalamus. Our resources and circuit map provide the common basis for the functional study of the basal ganglia in a small and optically tractable zebrafish brain for the comprehensive mechanistic understanding of the cortico-basal ganglia circuit.

## Introduction

The cortico-basal ganglia circuit plays crucial roles in learning and decision-making. The mammalian cortico-basal ganglia circuit consists of multiple brain regions widely distributed across the brain, which is too large to simultaneously image neuronal activities from the multiple interacting brain regions. One solution to this problem is using smaller teleost brains as an optically tractable miniature model of the vertebrate brain (Friedrich et al., 2010; Orger and De Polavieja, 2017; Vanwalleghem et al., 2018). Particularly, the telencephalon of adult zebrafish, where the main components of their evolutionary homolog of the mammalian cortico-basal ganglia circuit are located (Ganz et al., 2011; Mueller et al., 2008), is only about 1.5mm x 1.5mm in width and 1mm in depth, potentially providing by far a smaller model of the cortico-basal ganglia circuit. Previous studies have demonstrated that the dorsal part of the telencephalon is responsible for learning and decision-making both in the real world and virtual reality, which allows simultaneous calcium imaging of neuronal activities in the broad area of the telencephalon in awake-behaving zebrafish (Amo et al., 2014; Aoki et al., 2013; Frank et al., 2019; Huang et al., 2020; Lal et al., 2018; Torigoe et al., 2021).

In addition, zebrafish telencephalon shares the common basic structure with the mammalian forebrain. Teleost telencephalons have been known to undergo different developmental processes from those of mammals. While the mammalian neural tube evaginates, the dorsal part of the teleost neural tube, *i.e.*, the pallium, everts toward the outside, resulting in mediolaterally inverted topological correspondence between the mammalian and teleost telencephalons (Broglio et al., 2005; Northcutt and Braford, 1980) (Figure 1A). According to this scheme and previous anatomical studies, the mammalian cerebral cortex corresponds to the teleost pallium, while the mammalian striatum corresponds to the teleost dorsal (Vd) and central (Vc) nuclei of the ventral telencephalic area, which are located at the medial part of the subpallium (Figure 1B-C) (Mueller et al., 2008; 2011; Mueller and Wullimann, 2009; Rink and Wullimann, 2004; Wullimann and Mueller, 2004a). Consistently, the zebrafish pallium mainly consists of excitatory neurons with a small fraction of inhibitory neurons as in the case of the mammalian cerebral cortex, while the Vd/Vc region mostly consists of inhibitory neurons as in the case of the mammalian striatum (Aoki et al., 2013) (Figure S1A-E). Further anatomical mapping of the adult zebrafish telencephalon has been conducted mainly by chemical dye tracing for its connectivity (Rink and Wullimann, 2004; 2002; 2001; Yáñez et al., 2022), or by *in situ* hybridization (ISH) assay for the expression of genetic markers (Ganz et al., 2011; 2014; Mueller and Guo, 2009). These studies have elucidated detailed subdivisions of the zebrafish telencephalon and the evolutionary conservation of its overall framework with other vertebrates, as predicted from the developmental studies.

**Figure 1.**
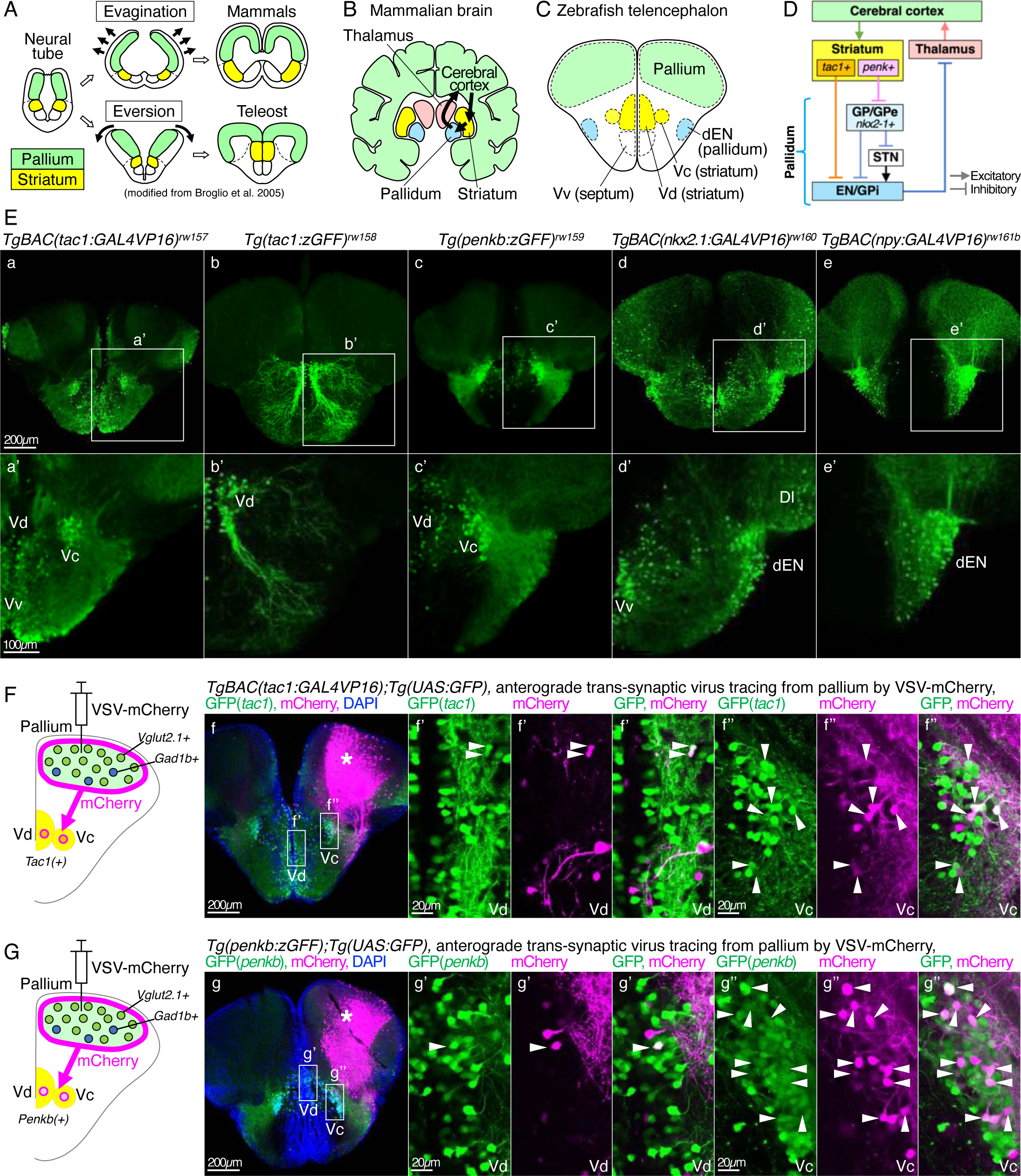
Transgenic and viral tools for anatomical characterization of specific neural subpopulations in the zebrafish basal ganglia. (A) Comparison of developmental processes of the mammalian and teleost brains (modified from (Broglio et al., 2005)). Areas colored with green and yellow indicate the pallium and the striatum, respectively. (B-C) Comparison of cortico-basal ganglia circuits of the mammalian and zebrafish brains. Coronal sections are shown. The evolutionarily homologous brain regions are colored in the same color. dEN, dorsal entopeduncular nucleus (homologous to the pallidum); Vv, ventral nuclei of the ventral telencephalic area (homologous to the septum). (D) Schematic diagram of the mammalian cortico-basal ganglia circuit. (E) Expression patterns of the newly established *Gal4*-driver lines for direct-pathway striatal neurons (*tac1* promoter, panels a-b), indirect-pathway striatal neurons (*penkb* promoter, panel c), and pallidal neurons (*nkx2.1* and *npy* promoters, panels d-e). Coronal sections in the anterior telencephalon are shown. Each *Gal4*-driver line was combined with *Tg(UAS:GFP)* or *Tg(UAS:AcGFP-P2A-WGA)* to visualize the expression pattern. Insets in the upper panels a-e indicate magnified areas in the lower panels a’-e’. (F) Illustration of a coronal slice depicting VSV-mCherry injection into the pallium and anterogradely labeled *tac1*+ neurons in the Vd/Vc. The panel f shows actual results from VSV-mCherry injection into the pallium of *TgBAC(tac1:GAL4VP16);Tg(UAS:GFP)* fish and immunohistochemistry of GFP (green), mCherry (magenta), and DAPI (blue). An asterisk indicates an approximate injection point. Insets show the positions of the panels f’ (focusing on Vd) and f’’ (focusing on Vc), where arrowheads indicate GFP and mCherry double-positive neurons. (G) Same as E, but in *Tg(penkb:zGFF);Tg(UAS:GFP)* fish.

Despite these anatomical studies on the zebrafish telencephalon, circuit mechanisms of the zebrafish cortico-basal ganglia circuit have not been studied. This is largely because anatomical characterization of zebrafish forebrain subdivisions is still insufficient, particularly regarding their correspondence to the mammalian basal ganglia. For example, evolutionary homologs of the mammalian GP/GPe (rodent globus pallidus/primate globus pallidus externus) and EN/GPi (rodent entopeduncular nucleus/primate globus pallidus internus) have not been clearly identified in zebrafish or any other teleosts. Therefore, neither the striatal direct- and indirect-pathways nor the pallido-thalamo-cortical output pathways have been described in teleosts. Other related brain regions such as the motor thalamus, subthalamic nucleus, and their connectivity to basal ganglia are also unidentified. Such lack of basic anatomical information has hampered the study of functions of zebrafish basal ganglia with a strict one-to-one assignment of each zebrafish brain region to a corresponding component of the cortico-basal ganglia circuit in mammals (Figure 1D).

Nevertheless, it is highly plausible that zebrafish do have such a conserved basal ganglia circuit based on phylogeny. Lamprey, a common ancestral species of teleosts and mammals, has been shown to have a full set of the mammalian basal ganglia components and connectivity (Grillner and Robertson, 2016; 2015; Grillner et al., 2013), suggesting that teleosts may inherit the conserved circuit structures.

In this work, using a set of newly generated cell-type specific *Gal4*-driver lines, we characterized the anatomical circuit structure of the telencephalon for each component of the cortico-basal ganglia circuit in adult zebrafish. Trans-synaptic tracing by genetically encoded neural tracer revealed that *tac1*-positive (+) direct-pathway striatal neurons project to *npy*+ output neurons in the dorsal entopeduncular nucleus (dEN), whereas *penkb*+ indirect-pathway striatal neurons project to *nkx2.1*+ neurons in the lateral nucleus of the ventral telencephalic area (Vl). The *nkx2.1*+ neurons in the Vl in turn project to the dEN, suggesting that the Vl and dEN are evolutionary homologs of the mammalian GP/GPe and EN/GPi, respectively. Further single-cell transcriptomic analysis identified *crhb+* neurons as another type of pallidal output neurons in the dEN. To our surprise, *npy*+ dEN neurons directly projected to the pallium, while *crhb*+ dEN neurons projected to the motor thalamus as well as the pallium. These results indicate that zebrafish have a shortcut output pathway from the pallidum to the pallium in addition to the evolutionarily conserved pallido-thalamic tract. Our comprehensive mapping of each basal ganglia component in the zebrafish telencephalon provides a structural basis of the simple and small model of the cortico-basal ganglia circuit for future functional studies.

## Results

### Transgenic tools for anatomical analysis of the zebrafish basal ganglia

For a comprehensive neuroanatomical study of the zebrafish basal ganglia, we established a set of new *Gal4*-driver lines which drive gene expression by the promoters of *tac1* and *penk*, *TgBAC(tac1:GAL4VP16), Tg(tac1:zGFF)*, and *Tg(penkb:zGFF)* (Figure 1E, panels a-c; Table S1). *Tac1* and *penk*, mammalian direct- and indirect-pathway striatal projection neuron markers, respectively (Gerfen and Young, 1988), are also known to be expressed in different subpopulations in the zebrafish Vd (Aoki et al., 2013). The *TgBAC(tac1:GAL4VP16)* and *Tg(penkb:zGFF)* drove gene expression in the Vd/Vc (Figure 1E, panels a’ and c’), where *tac1* and *penkb* are expressed (Figure S1F-G). The *Tg(tac1:zGFF)* drove gene expression mainly in the Vd and mostly lacked expression in the Vc (Figure 1E, panel b’).

We further established *Gal4*-driver lines that express transgenes in the pallidum. Historically, zebrafish pallidum had been thought to be intermingled in the Vd (Reiner and Northcutt, 1992; Wullimann and Mueller, 2004a; 2004b; Wullimann and Rink, 2002). Further anatomical mapping using molecular markers has identified the dEN as a putative pallidum, based on its expression of *gad1b* and *nkx2.1* (Ganz et al., 2011; Mueller and Guo, 2009) (Figure S1D and H). The dEN has been also known to express *neuropeptide Y* (*npy*) in zebrafish and other teleosts (Castro et al., 2006; Söderberg et al., 2000; Turner et al., 2016) (Figure S1I). Therefore, we generated *TgBAC(nkx2.1:GAL4VP16)* and *TgBAC(npy:GAL4VP16)* (Figure 1E, panels d-e; Table S1) and confirmed that these transgenic lines drive gene expression in the dEN (Figure 1E, panels d’-e’). The *TgBAC(nkx2.1:GAL4VP16)* also drove gene expression in the other telencephalic areas such as the septum (Vv) and the lateral zone of the dorsal pallium (Dl) (Figure 1E, panel d’; see also Figure S1H).

### The pallial neurons project to the *tac1*+ and *penkb*+ striatal neurons

Using the newly generated transgenic lines, we traced the pallio-striatal projection as a starting point for analyzing the zebrafish neural circuit equivalent to the cortico-basal ganglia circuit of mammals (see Figure 1D). Although zebrafish have been shown to have a putative pallio-striatal projection from the pallium to the ventral telencephalic area (Aoki et al., 2013; Rink and Wullimann, 2004), it is unclear whether this projection targets *tac1*+ and *penkb*+ striatal neurons as in the case of mammals. To elucidate this, we injected VSV-mCherry anterograde virus tracer into the pallium of *TgBAC(tac1:GAL4VP16);Tg(UAS:GFP)* fish (Figure 1F). Vesicular stomatitis virus (VSV) is capable of anterograde trans-synaptic infection from an injected site to its projection target (Beier et al., 2011), and therefore we used it to examine striatal cell types that actually receive projection from the pallium. We found trans-synaptically labeled cells expressing mCherry among the GFP-positive *tac1*+ Vd neurons (Figure 1F, panel f’) and Vc neurons (Figure 1F, panel f’’), indicating that the pallial neurons project to the *tac1*+ Vd/Vc neurons. We also performed the same experiment in *Tg(penkb:zGFF);Tg(UAS:GFP)* fish and found trans-synaptically labeled cells in the *penkb*+ Vd/Vc neurons (Figure 1G), indicating that the pallial neurons project to both *tac1*+ and *penkb*+ striatal neurons in the Vd/Vc.

### The *tac1+* and *penkb+* striatal neurons project to the different lateral subpallial regions, *i.e.* dEN and Vl, respectively

We then examined the striato-pallidal projection of zebrafish by transgenically expressing wheat germ agglutinin (WGA) with our *Gal4*-driver lines for the direct-pathway striatal neuron marker *tac1* (Figure 2A-B) or the indirect-pathway striatal neuron marker *penkb* (Figure 2C-D). WGA is a genetically-encoded neural tracer that is trans-synaptically transferred in both anterograde and retrograde manners (Yoshihara, 2002; Yoshihara et al., 1999). We used *Tg(UAS:AcGFP-2A-WGA)*, where labeling of both AcGFP and WGA indicates starter cells and labeling of only WGA indicates traced cells (Takeuchi et al., 2015). To label the putative zebrafish pallidal region dEN, we used anti-NPY antibody (Castro et al., 2006; Söderberg et al., 2000; Turner et al., 2016).

**Figure 2.**
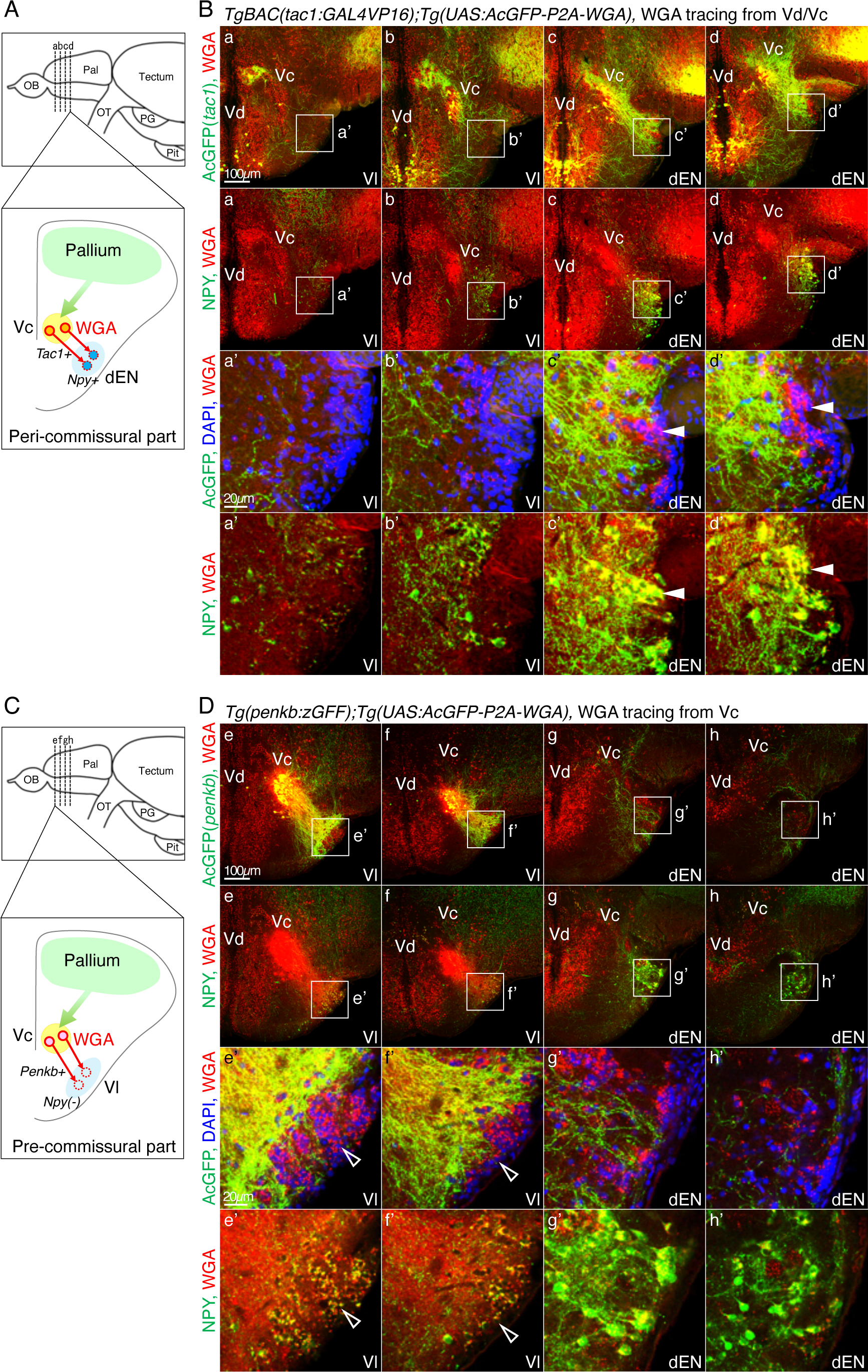
WGA tracing from the *tac1*+ and *penkb*+ striatal neurons caused WGA accumulation preferentially in the dEN and Vl, respectively. (A) Illustration of a coronal slice depicting WGA tracing from the *tac1*+ Vc neurons and trans-synaptic WGA transfer to the *npy*+ dEN neurons at the peri-commissural telencephalon. The top panel illustrates the lateral view of the zebrafish brain, and the dashed lines indicate antero-posterior positions of the coronal slices (a-d) shown in the panel B. OB, olfactory bulb; Pal, pallium; OT, optic tract; PG, preglomerular nucleus; Pit, pituitary. (B) WGA expression in the *tac1*+ Vd/Vc neurons and immunohistochemistry of AcGFP (green), WGA (red), DAPI (blue), and NPY (green). Four successive coronal slices (a-d) are shown so that the first two slices contain the NPY-negative Vl region and the latter two slices contain the NPY+ dEN region. Insets in the panels a-d (top two rows) show the positions of the panels a’-d’ (bottom two rows), focusing on the Vl or dEN. In the panels c’-d’, arrowheads indicate representative NPY+ dEN neurons with WGA signals. (C-D) Same as A-B, but from the *penkb*+ Vc neurons and trans-synaptic WGA transfer to the NPY-negative Vl neurons at the pre-commissural telencephalon. In the panels e’-f’, open arrowheads indicate representative NPY-negative Vl neurons with WGA signals.

In *TgBAC(tac1:GAL4VP16);Tg(UAS:AcGFP-2A-WGA)* fish, we found the *tac1*+ Vd/Vc neurons expressing AcGFP in the relatively posterior part of the telencephalon near the anterior commissure (peri-commissural part), and the AcGFP+ fibers mainly from the Vc innervated the NPY-immunoreactive neurons in the dEN (Figure 2B, panels c-d). We also found that the transgenic expression of WGA in the *tac1*+ Vd/Vc neurons caused a higher accumulation of WGA in the dEN (Figure 2B, panels c’-d’), in comparison to the anteriorly adjacent NPY-negative region Vl (Figure 2B, panels a’-b’). In contrast, in *Tg(penkb:zGFF);Tg(UAS:AcGFP-2A-WGA)* fish, AcGFP+ neurons located in the more anterior part of the Vc apart from the anterior commissure (pre-commissural part), and the AcGFP+ fibers innervated the Vl (Figure 2D, panels e-f). Consistently, the transgenic expression of WGA in the *penkb*+ Vc neurons caused a higher accumulation of WGA in the Vl (Figure 2D, panels e’-f’), in comparison to the dEN (Figure 2D, panels g’-h’). These results indicate that the *tac1*+ direct-pathway striatal neurons preferentially project to the dEN, whereas the *penkb*+ indirect-pathway striatal neurons preferentially project to the Vl. These results also suggest that the posterior part of the zebrafish pallidum dEN corresponds to the mammalian internal segment of the globus pallidus (EN/GPi), and the anterior part of the zebrafish pallidum Vl corresponds to the mammalian external segment of the globus pallidus (GP/GPe).

These projection patterns from the *tac1*+ and *penkb*+ striatal neurons seemed to be derived mostly from the Vc due to the weaker transgene expression in the Vd (Figure 2B and D, panels a-h). The contribution of the Vd was assessed by *Tg(tac1:zGFF);Tg(UAS:AcGFP-2A-WGA)*, which produced much higher expression of AcGFP and WGA in the Vd than that in the Vc (Figure S2A-B, panels a-e). The WGA expression mainly in the Vd caused WGA accumulation not only in the dEN (Figure S2B, panels c’-e’), but also in the ventral entopeduncular nucleus (vEN) and parvocellular preoptic nucleus anterior part (PPa) (Figure S2C). This result indicates that the Vd striatal subdivision projects to multiple target areas including the dEN whereas the Vc subdivision has a more specific projection to the dEN.

## The indirect pathway connectivity from the Vl to the dEN

We further transgenically expressed WGA by the *nkx2.1* promoter to identify downstream targets of the putative GP/GPe homolog Vl (Figure 3A). *Nkx2.1* is a developmental marker of pallidal neurons and is also an adult marker of the GP/GPe neurons in mammals (Abdi et al., 2015). We generated *TgBAC(nkx2.1:GAL4VP16);Tg(UAS:AcGFP-2A-WGA)* and found *nkx2.1*+ neurons expressing AcGFP in the Vl and the dEN (Figure 3B). Although a previous report did not observe *nkx2.1* expression in the Vl (Ganz et al., 2011), our ISH result showed a considerable number of *nkx2.1*+ neurons in the Vl (Figure 3C). These *nkx2.1*+ neurons did not overlap with NPY-immunoreactive neurons that were more preferentially distributed around the dEN (Figure 3B). Distribution of the *nkx2.1*+ neurons across the Vl-dEN region varied individually, some showed more *nkx2.1*+ neurons in the Vl (see Figure 3B) and the others showed more in the dEN (see Figure 3F). We further visualized WGA signals originating from the *nkx2.1+* neurons in the fish showing more *nkx2.1*+ neurons in the Vl and found WGA accumulation in the NPY+ dEN neurons (Figure 3D, filled arrowheads). This result is consistent with the evolutionary conservation of the indirect pathway in zebrafish. In mammals, the indirect pathway consists of two routes; the indirect projection via the subthalamic nucleus (STN) and the direct projection from the GP/GPe to the EN/GPi (Parent and Hazrati, 1995) (see Figure 1D). To distinguish whether the WGA transfer occurred directly from the Vl to the dEN or bi-synaptically through a zebrafish homolog of the STN, and to exclude the possibility of the WGA transfer from other surrounding *nkx2.1+* regions, we applied DiI in the Vl of the transgenic fish (Figure 3E-G). We confirmed that there were DiI and AcGFP double-positive fibers in the dEN, which originated from the *nkx2.1*+ neurons in the Vl, suggesting that there is a direct projection from the Vl to the dEN (Figure 3G).

**Figure 3.**
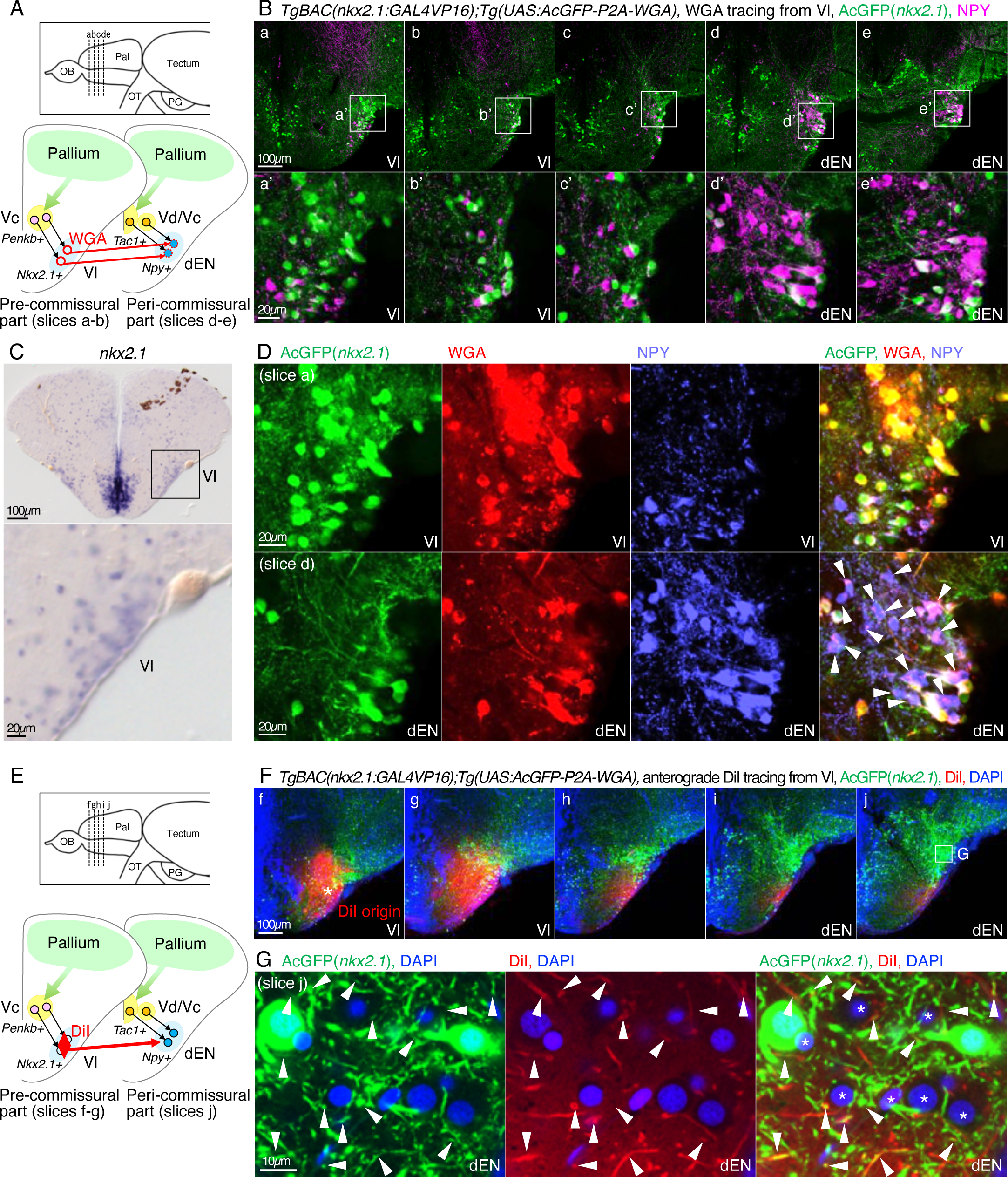
The indirect pathway targeted region Vl contains neurons which express GP/GPe marker *Nkx2.1* and project to the direct pathway targeted region dEN. (A) Illustration of coronal slices depicting WGA expression in the *nkx2.1*+ Vl neurons and trans-synaptic WGA transfer to the *npy*+ dEN neurons. (B) Different antero-posterior distributions of the *nkx2.1*+ neurons (AcGFP, green) and NPY+ neurons (NPY, magenta) in the five successive coronal slices (a-e). Insets in the upper panels a-e show the positions of the lower panels a’-e’, focusing on the Vl-dEN region. (C) ISH analysis of *nkx2.1* in the Vl. The same animal as the Figure S1H. (D) WGA tracing from the *nkx2.1*+ neurons and immunohistochemistry of AcGFP (green), WGA (red), and NPY (light blue). Upper panels show the Vl (slice a) and lower panels show the dEN (slice d). Arrowheads indicate NPY+ dEN neurons with WGA signals. (E) Illustration of coronal slices depicting anterograde DiI tracing from the Vl in *TgBAC(nkx2.1:GAL4VP16);Tg(UAS:AcGFP-P2A-WGA)* fish. (F) Five successive slices (f-j) of the DiI tracing from the Vl to the dEN, and multicolor imaging of AcGFP (green), DiI (red), and DAPI (blue). An inset in the panel e is magnified in G. (G) Magnified views of the dEN (slice j). Arrowheads indicate DiI and AcGFP double-positive fibers in the dEN, indicating direct projection from the *nkx2.1*+ Vl neurons to the dEN.

These results still do not exclude the possibility that there is an indirect pathway mediated by an STN, which has not been identified in zebrafish or other teleosts. In goldfish, it has been proposed that the nucleus ventromedialis thalami (VM), which positionally corresponds to the central posterior thalamic nucleus (CP) in zebrafish, is a possible subthalamic homolog (Yamamoto, 2009). Actually, the CP region contained neurons transgenically labeled by the promoter of the mammalian STN marker *pitx2* (D. M. Martin et al., 2004), suggesting the idea that the CP corresponds to the STN (Figure S3A, panel h; S3B). However, anterograde DiI tracing from the Vl did not stain any projection fiber around the CP (Figure S3C-D). DiI+ fibers that exited from the telencephalon were observed in the lateral forebrain bundle (LFB) without co-staining with AcGFP (Figure S3E), suggesting that the *nkx2.1+* neurons in the Vl mostly project intra-telencephalically. Neurons labeled by the *pitx2* promoter were also observed in the preglomerular nucleus (PG), but the PG seemed not to be an STN homolog because retrograde tracing from the PG by DiI or herpes simplex virus (HSV) did not label any Vl neurons (see Figure S4C and H, left panels).

### The *npy*+ dEN neurons directly project to the pallium

We then analyzed the basal ganglia output pathway from the dEN. First, we generated *TgBAC(npy:GAL4VP16);Tg(UAS:AcGFP-2A-WGA)* to express WGA in the *npy+* dEN neurons (Figure 4A). We found that the AcGFP+ axons from the *npy+* dEN neurons directly innervated the broad areas of the pallium (Figure 4B, panels a-g). WGA originating from the *npy+* dEN neurons was accumulated in most of the pallial neurons across multiple pallial subdivisions, including medial (Dm), central (Dc), and lateral (Dl) zone of the dorsal pallium (Figure 4C, panels d’-d’’’, arrowheads indicate representative pallial neurons with WGA signals). This result is consistent with previous observations of NPY-immunoreactive fibers in the pallium (Castro et al., 2006; Turner et al., 2016) and of retrogradely labeled cells in the dEN from various parts of the pallium by a chemical dye (Yáñez et al., 2022). The projection pattern was mostly intra-telencephalic and seemed not to exit to the diencephalic region that includes the thalamus (Figure 4B, panels h-n). The vEN, an adjacent pallidal structure projecting to the habenula (Amo et al., 2014), did not express AcGFP by the *npy* promoter (Figure 4D). We also confirmed that these NPY+ neurons in the dEN were GABAergic (Figure 4E), suggesting that these neurons have inhibitory roles on the pallium, as in the case of the mammalian EN/GPi neurons that control the cortex through the thalamus.

**Figure 4.**
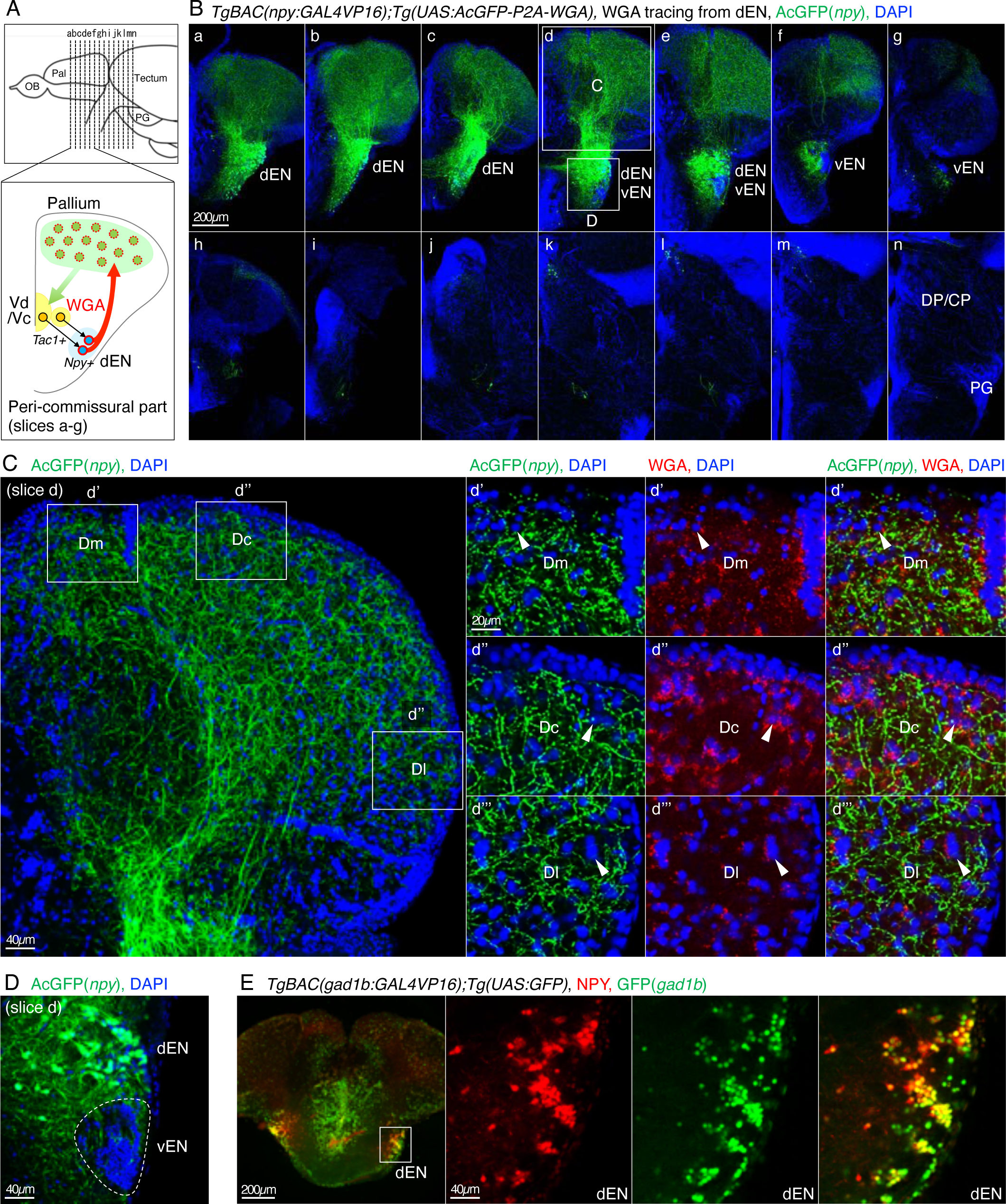
The *npy*-positive inhibitory neurons in the dEN directly project to the pallium, and not to the thalamus. (A) Illustration of a coronal slice depicting WGA expression in the *npy*+ dEN neurons and trans-synaptic WGA transfer to the pallial neurons. (B) Projection patterns of the *npy*+ neurons (AcGFP, green) to the pallium (DAPI, blue) in 12 successive coronal slices (a-n). In the slice d, insets at the pallium and the dEN-vEN region are magnified in C and D, respectively. (C) WGA tracing from the *npy*+ dEN neurons and immunohistochemistry of AcGFP (green), WGA (red), and DAPI (blue). The left panel shows a magnified view of the pallium in the slice d, and insets show the positions of the right panels d’-d’’’, focusing on the different pallial subdivisions Dm, Dc, and Dl. In the panels d’-d’’’, arrowheads indicate representative pallial neurons with WGA signals. (D) A magnified view of the dEN-vEN region in the slice d. The vEN is circled with a dotted line. (E) Immunohistochemistry of *gad1b*+ neurons (GFP, green) and *npy*+ neurons (NPY, red) in *TgBAC(gad1b:GAL4VP16);Tg(UAS:GFP)* fish. An inset in the leftmost panel is magnified in the right three panels. NPY immunoreactive cells were co-stained with GFP signals (rightmost panel).

### The *npy*-negative dEN neurons project to the thalamus

Then, we asked whether there are any dEN neurons that project to the thalamus in zebrafish. We performed retrograde tracing of pallido-thalamic output neurons from the dorsal posterior thalamic nucleus (DP) and the CP, as a candidate region of the motor thalamus in zebrafish. The DP/CP is located in the medial diencephalic area similar to the mammalian thalamus, and receives several sensory inputs directly or indirectly (Echteler and Saidel, 1981; Goodson and Bass, 2002; Mueller, 2012; Northcutt, 2006). DiI application to the DP/CP caused retrograde DiI labeling of *gad1b*-positive dEN neurons (Figure 5A, filled arrowhead; Table S2), indicating that there is an inhibitory neural subpopulation projecting to the DP/CP. In addition, the retrogradely labeled DP/CP projecting dEN neurons were *npy*-negative (Figure 5B, open arrowheads; Table S2), consistently with the intra-telencephalic projection pattern of the *npy*+ dEN neurons (see Figure 4B).

**Figure 5.**
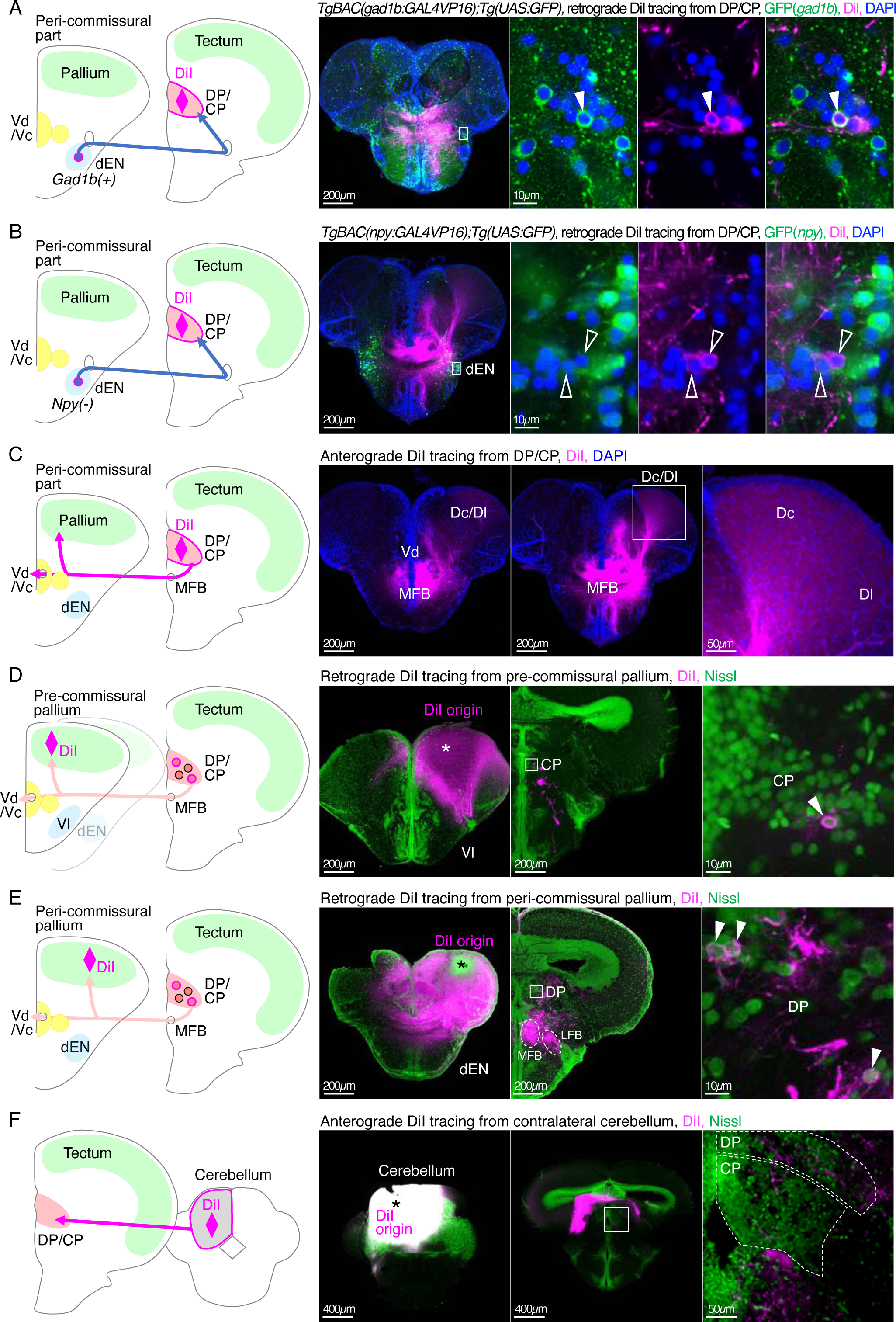
The *npy*-negative inhibitory neurons in the dEN project to the DP/CP thalamus and form the pallido-thalamo-pallial pathway. (A) DiI tracing from the DP/CP in *TgBAC(gad1b:GAL4VP16);Tg(UAS:GFP)* fish retrogradely labeled *gad1b*+ neurons in the dEN. An inset in the leftmost panel is magnified in the right three panels. Filled arrowheads indicate retrogradely labeled *gad1b*+ neurons in the dEN. (B) Same as A, but in *TgBAC(npy:GAL4VP16);Tg(UAS:GFP)* fish. Open arrowheads indicate retrogradely labeled *npy*-negative dEN neurons. (C) DiI tracing from the DP/CP anterogradely labeled projection fibers in the MFB and the Dc/Dl pallium. The same animal as panel B. Two successive peri-commissural slices are shown. An inset in the middle panel is magnified in the rightmost panel. (D) DiI tracing from the pre-commissural pallium retrogradely labeled thalamic neurons. An asterisk in the left panel indicates DiI origin. An inset in the middle panel is magnified in the rightmost panel. An arrowhead in the rightmost panel indicates retrogradely labeled CP neurons. (E) Same as D, but from the peri-commissural pallium. Arrowheads in the rightmost panel indicate retrogradely labeled DP neurons. (F) DiI tracing from the cerebellum anterogradely labeled projection fibers in the contralateral DP/CP. An asterisk in the left panel indicates DiI origin. An inset in the middle panel is magnified in the rightmost panel.

We then tested if the basal ganglia-recipient thalamic region DP/CP has projection back to the pallium. DiI application to the DP/CP caused anterograde labeling of projection fibers in the pallium (Figure 5C). In addition, DiI application to the pre- and peri-commissural pallium retrogradely labeled neurons in the DP/CP (Figure 5D-E, filled arrowheads), indicating that the DP/CP has projection to the pallium and can form a loop circuit equivalent to the cortico-basal ganglia-thalamo-cortical loop of mammals. We also found that DiI application to the cerebellum anterogradely labeled projection fibers from the cerebellum to the DP/CP (Figure 5F), supporting our idea that the DP/CP is a teleost counterpart of the mammalian motor thalamus (Middleton and Strick, 2000).

We also performed retrograde tracing from the PG, which is another counterpart of the mammalian thalamus. The PG is a midbrain-derived thalamic-like structure that receives various sensory inputs and has strong projection to the pallium (Bloch et al., 2020; Mueller, 2012; Yamamoto and Ito, 2008; 2005). We applied DiI to the PG (Figure S4A-B) and found no retrogradely labeled neuron in the dEN (Figure S4C), indicating that the PG is not innervated by the dEN. Instead, we found anterogradely labeled projection fibers and retrogradely labeled neurons in the pallium (Figure S4D-E). Retrograde virus tracing from the PG by HSV-GFP also resulted in retrograde labeling not in the dEN but in the pallium (Figure S4F-J). These results indicate that the PG does not receive input from the dEN but has reciprocal connections with the pallium, and may suggest its dedicated role for processing sensory information.

### ScRNAseq analysis identified *crhb* as a candidate marker of the thalamus-projecting dEN neurons

To identify genetic markers of the *npy*-negative thalamus-projecting neurons in the dEN, we conducted single-cell RNA sequencing (scRNAseq) analysis. We dissected the dEN and its surrounding regions from six adult individuals of *TgBAC(npy:GAL4VP16);Tg(UAS:GFP)* (Figure 6A and S5A). We then dissociated the dissected tissue and used the droplet-based 3’end scRNAseq system Chromium (10x Genomics). We obtained transcriptomic data from 3,381 cells and performed unbiased clustering by Seurat (Butler et al., 2018) (see STAR Methods for the details). We found 18 clusters including glia and hemocytes, and 12 clusters were neuronal clusters that expressed neuronal markers *snap25, sv2a,* and *sty1* (Figure S5B-C and Data S1) (Cosacak et al., 2019; A. Martin et al., 2022). Among the neuronal clusters, six clusters were identified as entopeduncular clusters due to their expression of GABAergic neuronal markers (*gad1b, gad2,* and *slc17a7a*), pallidum developmental precursor markers (*nkx2.1* and *lhx6*), and zebrafish dEN marker *npy* (EN1 to EN6 clusters, Figure 6B and S5D-I). Another GABAergic cluster was striatal neurons expressing *penkb* (Vc cluster, Figure S5J-K). In mammals, the EN/GPi expresses *parvalbumin (pv)* in thalamus-projecting neurons and *somatostatin (sst)* in habenula-projecting neurons (Wallace et al., 2017). In zebrafish, all of the EN clusters were negative for *pv* and positive for *sst*, hence the *pv*/*sst* axis was not useful to distinguish subpopulations in the EN clusters (Figure S5L-P). Among the other remaining neuronal clusters, four clusters were pallial excitatory neurons that expressed glutamatergic neuronal markers (*slc17a6b* and *slc17a6a*) and pallium markers (*neurod1 and emx1*) (Pallium 1 to 4 clusters, Figure S5Q-T) (Ganz et al., 2011). Another glutamatergic cluster was identified as a vEN cluster due to its expression of *lhx5* (vEN cluster, Figure S5U) (Turner et al., 2016).

**Figure 6.**
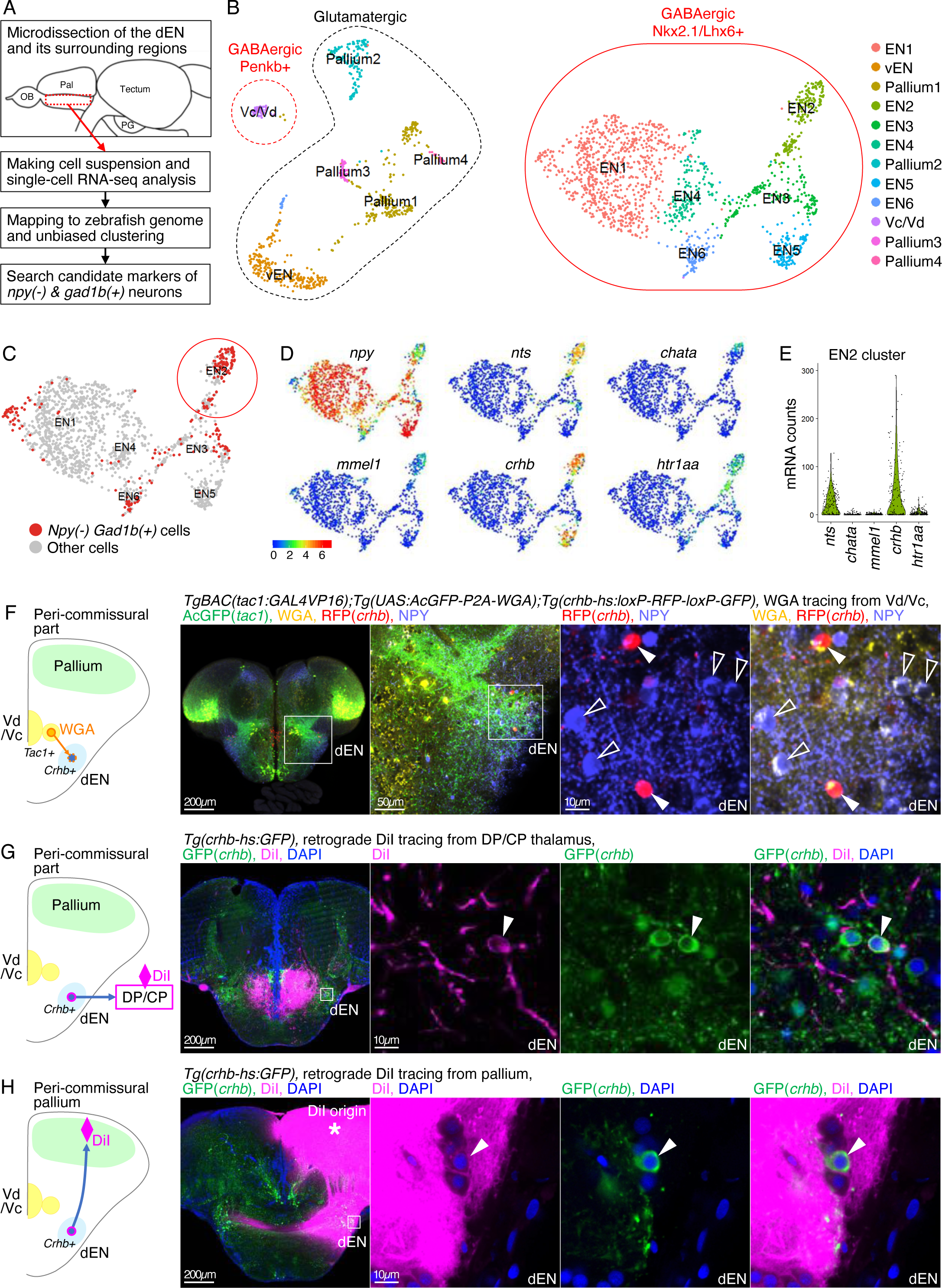
ScRNAseq analysis of the zebrafish pallidum identified *crhb* as a marker of the thalamus-projecting dEN neurons, which also directly project to the pallium. (A) An overview of the experimental strategy of the scRNAseq analysis. (B) UMAP plot displaying the result of clustering. Each dot represents one cell and clusters are color-coded. Only neuronal clusters are shown. Clusters intrinsic to entopeduncular neurons are circled with the solid red line (EN1 to 6 clusters). (C) UMAP plot showing the distribution of the [*npy*-negative and *gad1b*-positive] cells (red dots) in the six EN clusters. These cells highly accumulated in the EN2 cluster (red circle). (D and E) Expression patterns of the top-five differentially expressed genes of the EN2 cluster*. Crhb* showed the highest expression among them. (F) WGA tracing from the *tac1*+ Vd/Vc neurons and trans-synaptic WGA transfer to *crhb*+ dEN neurons. Immunohistochemistry of AcGFP (green), WGA (yellow), *crhb* (magenta), and NPY (light blue). Insets show the positions of the next panels, focusing on the dEN. Filled arrowheads indicate *crhb*-positive and *npy*-negative neurons that accumulate WGA as much as that of *crhb*-negative and *npy*-positive neurons (open arrowheads). (G) DiI tracing from the DP/CP in *Tg(crhb-hs:GFP)* fish retrogradely labeled *crhb*+ dEN neurons. An inset in the leftmost panel is magnified in the right three panels. Arrowheads indicate retrogradely labeled *crhb*+ dEN neurons. (H) Same as G, but from the pallium. An asterisk indicates DiI origin.

We further defined [*npy*-negative and *gad1b*-positive] cells, which are characteristics of the thalamus-projecting dEN neurons (see Figure 5A-B), based on expression levels of the *npy* and *gad1b* mRNA (Figure S5V-W). Among the six EN clusters, EN1, EN2, EN3, and EN6 clusters had significant amount of [*npy*-negative and *gad1b*-positive] cells (Figure 6C), and the largest number (98 cells) and coverage (53.8%) of such cells were found in the EN2 cluster (Table S3). The top-five differentially expressed genes of this [*npy*-negative and *gad1b*-positive] cells in the EN2 cluster were *nts*, *chata*, *mmel1*, *crhb*, and *htraa1* (Data S2). Among these, *crhb* had the highest expression level in the EN2 cluster (Figure 6D-E), and therefore we decided to further analyze *crhb* as a candidate marker of the thalamus-projecting dEN neurons. We then performed two-color fluorescent ISH of *npy* and *crhb* (Figure S6A), and found three types of neurons that are consistent with the scRNAseq result: (1) *npy* only-positive neurons (seen in the EN1, 3, 4, 6 clusters), (2) *npy* and *crhb* double-positive neurons (seen in the EN5 cluster, filled arrowheads in Figure S6A), (3) *crhb* only-positive neurons (seen in the EN2 cluster, open arrowheads in Figure S6A).

### The *crhb*+ dEN neurons project to the thalamus as well as directly to the pallium

We established *Tg(crhb-hs:loxP-RFP-loxP-GFP)* and *Tg(crhb-hs:GFP)* to transgenically label *crhb*+ neurons (Figure S6B-C). We confirmed that the *crhb* only-positive dEN neurons receive projection from the direct-pathway striatal neurons in *TgBAC(tac1:GAL4VP16);Tg(UAS:AcGFP-2A-WGA);Tg(crhb-hs:loxP-RFP-loxP-GFP)* fish: the transgenic expression of WGA in the *tac1*+ Vd/Vc neurons caused accumulation of WGA in the *crhb*-positive and NPY-negative neurons in the dEN (Figure 6F). DiI application to the DP/CP in the *Tg(crhb-hs:GFP)* fish caused retrograde DiI labeling of the *crhb*-positive dEN neurons (Figure 6G, filled arrowhead; Table S2). This result, in combination with the absence of *npy* expression in the DP/CP projecting dEN neurons (see Figure 5B), indicates that the *crhb* only-positive dEN neurons project to the DP/CP. In addition, detailed observation of the *Tg(crhb-hs:GFP)* fish suggested that the *crhb*-positive dEN neurons may also project to the pallium (Figure S6D, filled arrows). To our surprise, we actually found that DiI application to the pallium caused retrograde DiI labeling of the *crhb*-positive dEN neurons (Figure 6H). These results indicate that there is also a direct projection from the *crhb*-positive dEN neurons to the pallium as well as the projection to the DP/CP thalamus.

### The motor output pathway derived from the pallium

Lastly, we searched for a motor output pathway from the pallium. Although teleosts are thought to lack a direct projection from the pallium to the spinal cord (Yamamoto et al., 2017), it is still possible that there are functional counterparts of the mammalian corticospinal tract, which mediate motor commands from the pallium to the downstream motor circuits. In teleosts, the optic tectum, which is homologous to the mammalian superior colliculus, is known to control body movements (Helmbrecht et al., 2018; Isa et al., 2021). Therefore, to test whether the pallium has a projection to the optic tectum, we visualized output pathways from the pallium using VSV-mCherry anterograde trans-synaptic virus tracer. Injection of VSV-mCherry into the pallium caused labeling of the virus-infected cells in various brain regions (Figure 7A-B and S7A-C), including the optic tectum (Figure 7B, panel p). The VSV-infected tectal cells were mainly located in the stratum periventriculare (SPV) (a.k.a. periventricular gray zone) of the ipsilateral and contralateral tectum (Figure 7C-D). In the SPV, the shallower layers contained VSV-infected neurons, putatively periventricular projection neurons or interneurons based on its location (Nevin et al., 2010). The deeper layers contained VSV-infected cells with “bottlebrush-like” morphology extending to the pial surface (Figure S7D), indicating that these cells are radial glia (or radial astrocytes) (Sild et al., 2016; Xiao et al., 2011). Tectal radial glia are known to receive retinal synaptic input in flogs (Benfey et al., 2021) and might contribute to sensorimotor control as shown in the larval hindbrain circuit (Mu et al., 2019). Injection of VSV-mCherry into the pallium of another fish visualized entering pallio-tectal projection fibers from the LFB to the ipsilateral tectum (Figure 7E and S8A-F). We also found virus-infected cells in the contralateral pallium (Figure S7E), Vd/Vc (Figure S7F-G), DP/CP (Figure S7H), and PG (Figure S7I), and mCherry+ projection fibers from the injected pallium in the caudal zone of the periventricular hypothalamus (Hc) (Figure S7J). We further confirmed the pallio-tectal projection by retrograde virus tracing from the tectum using HSV-GFP (Figure 7F). We found several retrogradely labeled pallial neurons mainly in the ipsilateral pallium (Figure 7G-J) although only a small fraction of the tectum was subject to the injection (Figure 7K). This result indicates that there is a direct projection from the pallium to the tectum, which could be a pallial motor output pathway of teleosts. This possibility has also been previously proposed in the goldfish studies (Meek, 1983).

**Figure 7.**
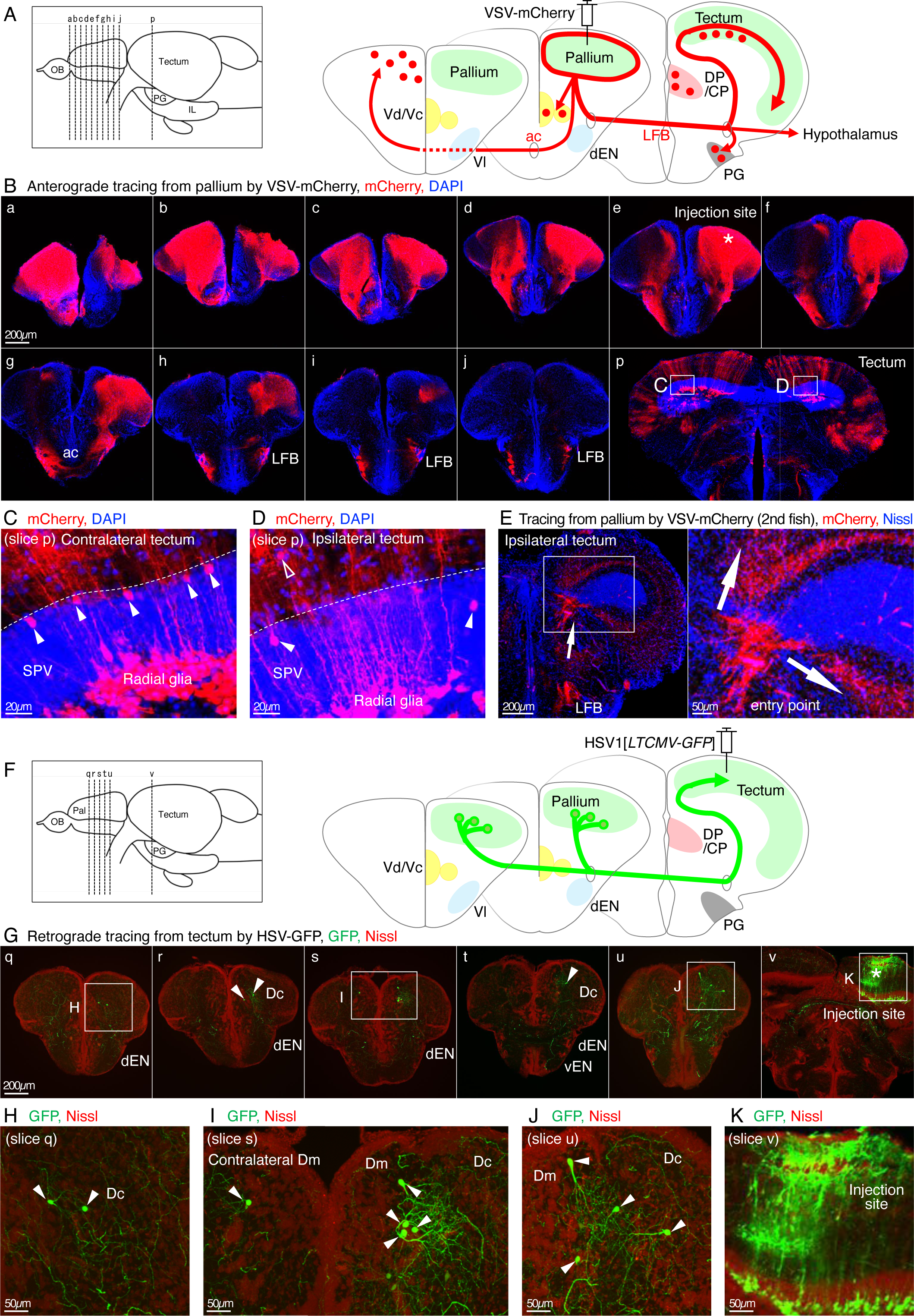
Visualization of output pathways from the pallium to the optic tectum. (A) Illustration of coronal slices depicting VSV-mCherry injection into the pallium and distribution of anterogradely trans-synaptically labeled neurons. (B) VSV-mCherry injection into the pallium and immunohistochemistry of mCherry (red), and DAPI (blue). Ten successive coronal slices at the telencephalon (a-j) and one slice at the tectum (p) are shown. An asterisk in the panel e indicates an approximate injection point. Insets at the left and right tectum in the slice p are magnified in C and D, respectively. (C and D) Magnified views of the bilateral tectum in the slice p. Dotted line indicates the boundary between the SPV and the upper layer. Filled arrowheads indicate labeled tectal neurons in the shallower layers of the SPV, and an open arrowhead indicates one outside the SPV. Radial glia were also labeled in the deeper layers of the SPV. (E) VSV-mCherry injection into the pallium of another fish visualized entering pallio-tectal projection fibers from the LFB to the ipsilateral tectum (arrows). An inset is magnified in the right panel. (F) Illustration of coronal slices depicting HSV-GFP injection into the tectum and retrogradely labeled neurons in the pallium. (G) HSV-GFP injection into the tectum and immunohistochemistry of GFP (green) and Nissl (red). Five successive coronal slices at the telencephalon (q-u) and one slice at the tectum (v) are shown. Arrowheads indicate retrogradely labeled pallial neurons. Insets show the positions of the panels H-J (focusing on the pallium) and K (focusing on the injection site in the tectum). An asterisk indicates an approximate injection point. (H-J) Magnified views of the retrogradely labeled pallial neurons (arrowheads). (K) A magnified view of the injection site in the tectum.

**Figure 8.**
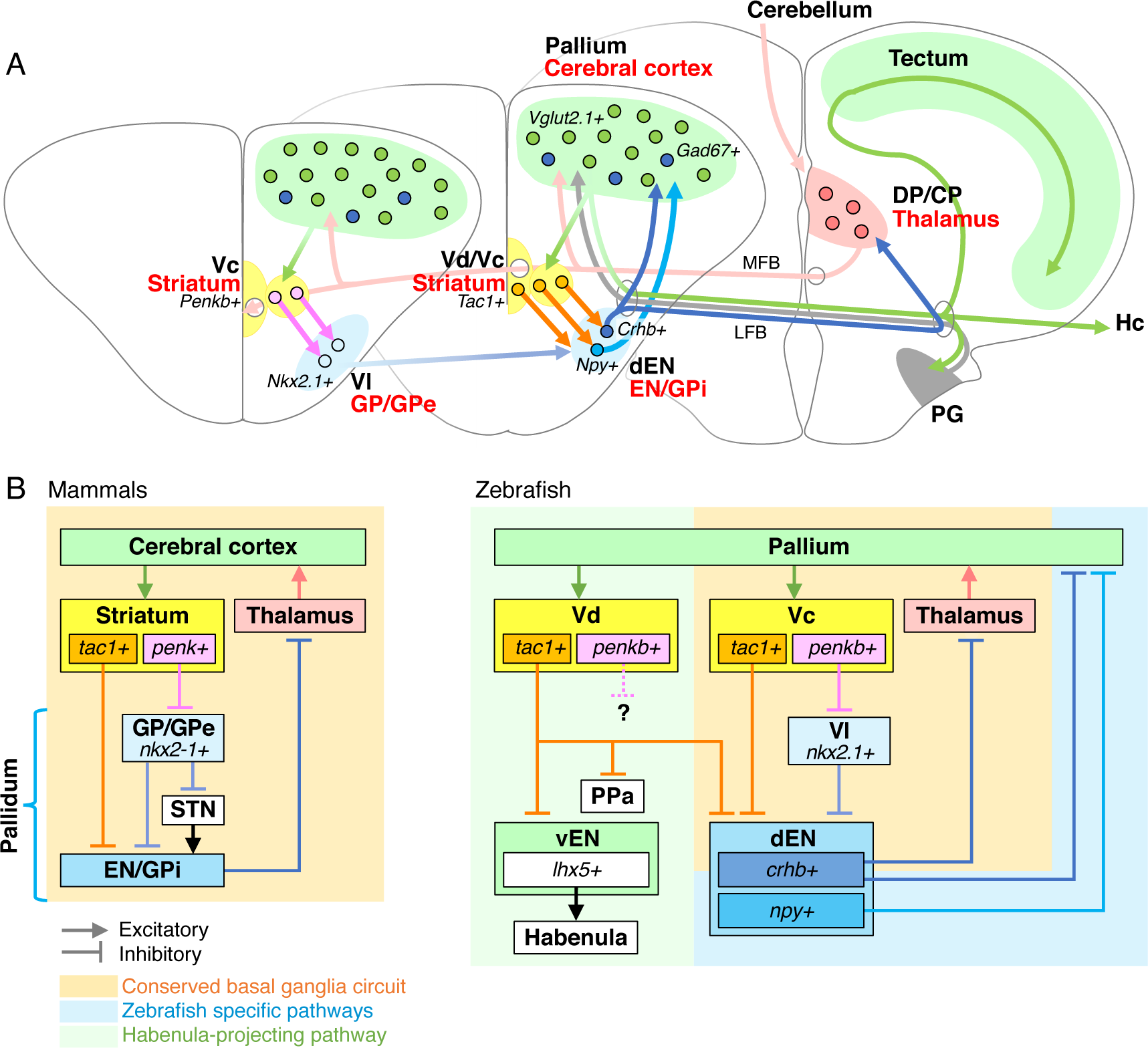
The anatomical map of the zebrafish cortico-basal ganglia circuit. (A) A schematic diagram of the zebrafish cortico-basal ganglia-thalamic network elucidated in this study. (B) Comparison between the mammalian and zebrafish cortico-basal ganglia circuits. The conserved parts are colored with orange. The zebrafish-specific part is colored with blue. The habenula-projecting pathway of zebrafish is colored with green.

## Discussion

In this study, we made a comprehensive anatomical map of the adult zebrafish cortico-basal ganglia circuit, in a manner that is highly comparable to that of mammals. Adult zebrafish display a full repertoire of mature behaviors, such as social conflict behavior, active avoidance, reinforcement learning, and decision-making (Amo et al., 2014; Aoki et al., 2013; Cherng et al., 2020; Chou et al., 2016; Frank et al., 2019; Huang et al., 2020; Kalueff et al., 2013; Lal et al., 2018; Orger and De Polavieja, 2017). In addition, recent methodological advances allowed two-photon calcium imaging of telencephalic neuronal activity in head-fixed adult zebrafish during learning and decision-making in a virtual reality environment (Huang et al., 2020; Torigoe et al., 2021). The telencephalon of adult zebrafish is only about 1.5mm x 1.5mm wide and 1mm deep, and thus broad areas of the pallium, striatum, and pallidum can be simultaneously recorded by calcium imaging and/or manipulated by optogenetics based on two-photon laser scanning. Furthermore, three-photon laser scanning allows functional imaging at cellular resolution throughout the telencephalon in intact adult zebrafish (Chow et al., 2020). Our anatomical map will allow one-to-one assignment of such imaging/optogenetics results from the zebrafish basal ganglia on the common basis of the mammalian basal ganglia studies for comprehensive mechanistic understanding of the cortico-basal ganglia circuit.

### The conserved direct and indirect pathways in the zebrafish basal ganglia

We revealed that there are conserved direct- and indirect-pathways originating from the distinct striatal subpopulations in teleosts. The *tac1*+ (direct-pathway) and *penkb*+ (indirect-pathway) striatal neurons projected to different pallidal subregions dEN and Vl, respectively. Based on this, the dEN and Vl were identified as the evolutionary homologs of mammalian EN/GPi and GP/GPe, respectively. Consistently, the GP/GPe corresponding region Vl expressed *nkx2.1*, which is also known as a mammalian GP/GPe marker. It has been shown that the basic structure of the basal ganglia had already existed in the lamprey, the most ancient group of vertebrates (Grillner et al., 2013; Grillner and Robertson, 2016; 2015). Our study revealed that most of the basic components and connectivity of the cortico-basal ganglia circuit are also shared in zebrafish.

In zebrafish, the two pallidal subnuclei Vl and dEN were segregated antero-posteriorly, in contrast to the mammalian GP/GPe and EN/GPi, which are segregated medio-laterally. This segregation is surprising because pallidal subpopulations corresponding to the mammalian GP/GPe and EN/GPi are thought to be intermingled in a single pallidal nucleus in lampreys, amphibians, reptiles, and birds (Parra et al., 1998; SMEETS et al., 2000; Stephenson-Jones et al., 2011). Our result suggests that segregation of the pallidal subregions has occurred multiple times in the vertebrate phylogeny; one is early in the mammalian lineage (Reiner et al., 1998) and the other is somewhere in the actinopterygian lineage.

### The zebrafish-specific direct output pathways from the pallidum to the pallium

In contrast to the striato-pallidal pathways, the zebrafish pallidal output pathways appear to have unique characteristics compared with those of mammals. In addition to the conserved pallido-thalamo-pallial output pathway from the dEN, we identified the fish-specific shortcut output pathway which directly projects from the *npy*+ dEN neurons to the pallium without mediating the thalamus. This pathway is unignorably robust: the *npy*+ dEN neurons had highly elaborated axon terminals in the pallium, which caused accumulation of WGA in almost all the cells in the projected area (see Figure 4C). We also found that *crhb*+ neurons in the dEN, which compose the conserved pallido-thalamo-pallial output pathway, also directly project to the pallium. In these zebrafish pallidal output neurons, genetic markers are also divergent; neither *npy* nor *crhb* is known to be expressed in the mammalian pallidum (except that the ventral pallidum contains *npy*+ neurons in rodents (Zaborszky et al., 2012)).

The existence of NPY-immunoreactive cells in the lateral subpallium has been reported in many other teleosts (Castro et al., 1999; García-Fernández et al., 1992; Kawaguchi et al., 2018; Pontet et al., 1989; Saha et al., 2014; Traverso et al., 2003) and white sturgeon (Chiba and Honma, 1994). Projection from the lateral subpallium to the pallium has also been reported in several teleosts (Folgueira et al., 2004; Murakami et al., 1983; Northcutt, 2006). In Polypterus, however, the lateral subpallium contained neither NPY immunoreactive cell nor retrogradely labeled cell from the pallium (Holmes and Northcutt, 2003; Reiner and Northcutt, 1992), suggesting that the direct output pathway from the pallidum to the pallium has been acquired during the evolution of actinopteri.

### The direct projection from the Vl to the dEN without mediating an STN

In this study, we were not able to identify projections from the Vl to an STN in zebrafish. Moreover, the STN itself has not been identified in zebrafish or other teleosts. In goldfish, it has been hypothesized that the VM, nucleus ventromedialis thalami, is a possible STN homolog (Yamamoto, 2009). Positionally, this VM region of goldfish corresponds to the CP of zebrafish, which has been already characterized as one of the basal ganglia-recipient thalamic regions in this study based on its connection from the dEN and to the pallium. Thus, there may be a STN homolog in the unexplored brain areas in this study. A bold hypothesis is that zebrafish lack an STN. Even in this case, the functional connectivity of the indirect-pathway can be accomplished by the direct projection from the Vl to the dEN (the STN is excitatory and its presence or absence does not affect the excitatory or inhibitory nature of the pathway’s net transmission).

What is the meaning of the missing pallidal projection to the subthalamus? Together with the aforementioned fish-specific direct output pathways from the dEN to the pallium, there may be an evolutionary trend of enhanced intra-telencephalic connection that substitutes diencephalon-mediated pathways. In the fish brain, the telencephalon is located at the anterior end and there is significant constriction between the telencephalon and the diencephalon. Because of this constriction, the geometrical distance between the telencephalon and the diencephalon becomes longer, and projection back and forth between them can be developmentally costly. In addition, the constriction can be a bottleneck to information transfer due to its small cross-sectional area. This situation may have driven evolution so that the intra-telencephalic connectivity is enhanced in the teleost forebrain to substitute possibly compromised functions of the thalamus- and subthalamus-mediated pathways. To summarize, our results revealed not only the evolutionary conservation but also the specialization of the cortico-basal ganglia circuit of zebrafish in comparison with those of other vertebrates.

### Application of the transgenic zebrafish lines for functional analyses of the basal ganglia

Besides the anatomical characterization of the circuit, our *Gal4*-driver lines have by far more versatile use to elucidate the basic mechanisms of the basal ganglia. For example, we have recently shown that the zebrafish pallium is capable of generating learning-dependent predictive coding for the efficient control of adaptive escape behavior (Torigoe et al., 2021). To know how prediction and prediction error can be represented by the pallial neurons and how they are used for the control of behavior, it is essential to analyze detailed interactions between the pallium and the basal ganglia at the cellular and region-wide levels. Our transgenic lines can be used for such studies by expressing genetically-encoded calcium indicators/channelrhodopsins in the anatomically-defined striatal and pallidal neurons. For example, it is possible to perform simultaneous two-photon calcium imaging of the pallium and the striatum which are only ∼500 µm apart in depth. It is also possible to analyze learning-dependent cortico-striatal plasticity, by imaging the striatal responses to optogenetic manipulation of pallial neurons by holographic light focusing (Carrillo-Reid et al., 2016; Fisek et al., 2023; Marshel et al., 2019).

In mammals, interconnected cortical and striatal regions through the cortico-basal ganglia loop are far more distantly apart compared with zebrafish and simultaneous cellular-resolution imaging/manipulation of such a wide area is technically difficult. Zebrafish offer opportunities to analyze detailed interactions between multiple components of the small cortico-basal ganglia circuit for elucidation of circuit- and cellular-level implementation of the complex computational functions underlying learning and decision-making.

## Author contributions

H.O. conceived and supervised the project. Y.T., H.K., and H.O. designed the experiments and analyses. Y.T. and H.K. carried out the experiments and analyses. Y.T., R.A., T.S., and S.H. established transgenic lines. H.K. established VSV and HSV virus tracing systems. Y.T. and H.K. carried out the scRNAseq experiments and analyses. Y.T., H.K., and H.O. wrote the manuscript.

## Supporting information

Tanimoto_et_al_Supplementary_Information

## Acknowledgments

We thank all of the members of our laboratory for the discussions and the fish care support, and the RIKEN Research Resource Division for supporting experimental devices. We also thank Drs. M. Miyasaka and Y. Yoshihara for providing the *Tg(UAS:AcGFP-2A-WGA)* line, and Dr. K. Kawakami for providing the *Tg(UAS:GFP)* line and the *zGFF* construct. We also thank Drs. N Yamamoto and H Hagio for their advice on the interpretation of the fish anatomical data. This work was supported by JSPS KAKENHI Grant-in-Aid for Scientific Research (23H04976, 22H05520), Grant-in-Aid for Early-Career Scientists (JP20K15942), and was partially supported by RIKEN-Kao Collaboration Center.

## STAR★Methods

### KEY RESOURCES TABLE

#### RESOURCE AVAILABILITY

##### Lead Contact

Further information and requests for resources and reagents should be directed to and will be fulfilled by the Lead Contact, Hitoshi Okamoto.

##### Materials Availability

All the transgenic lines, plasmids, and virus vectors generated in this study are available from the lead contact upon request. See the Key Resources Table for the details.

##### Data and Code Availability

ScRNAseq data and R scripts used for the data analysis have been deposited at GEO and are publicly available as of the date of publication. Accession numbers are listed in the key resources table. Microscopy data reported in this paper will be shared by the lead contact upon request. Any additional information required to reanalyze the data reported in this paper is available from the lead contact upon request.

### EXPERIMENTAL MODEL AND SUBJECT DETAILS

All protocols were reviewed and approved by the Animal Care and Use Committees of the RIKEN Center for Brain Science. This study used adult wild-type zebrafish (RIKEN-Wako, Saitama, Japan) and transgenic zebrafish lines. Transgenic lines generated in this study are shown in Table S1. The transgenic lines published previously and applied in this study are the following: The *Tg(UAS:GFP)* (Asakawa et al., 2008), *Tg(UAS:AcGFP-P2A-WGA)* (Takeuchi et al., 2015). All the transgenic zebrafish were maintained and selected by their fluorescence patterns in the larva stage. Fish were maintained in 7-L tanks with continuous water exchange at 28.5°C under the 14-hour light/10-hour dark cycling.

### METHOD DETAILS

#### Generation of transgenic animals

For the transgenic lines generated by transposon-mediated BAC, we used *tac1, nkx2.1, npy,* and *gad1b* containing BAC clones zC135B21, zC62I6, zC72N5, and zC68B9 to establish *TgBAC(tac1:GAL4VP16), TgBAC(nkx2.1:GAL4VP16), TgBAC(npy:GAL4VP16),* and *TgBAC(gad1b:GAL4VP16)*, respectively, as shown in our previous report (Amo et al., 2014). Briefly, each BAC clone was identified by blasting cDNA sequences of each gene against zebrafish genomic sequences of Ensemble (http://uswest.ensembl.org/Danio_rerio/Info/Index), which provides the BAC end information. The BAC plasmids containing the *GAL4VP16* sequence (Agetsuma et al., 2010) were generated by homologous recombination as described previously and used for creating the transgenic fish (Kimura et al., 2006). To facilitate genomic integration of the BAC constructs, the *iTol2* cassette was introduced by homologous recombination in bacteria (Suster et al., 2009). The purified plasmids were injected into 1-2 cell stage zebrafish eggs, and then these injected embryos were grown for further fluorescent screening of next-generation larvae by crossing with the *Tg(UAS:EGFP)* fish.

For the transgenic lines generated by CRISPER/Cas9 knock-in strategy, we used *zGFF*, a modified version of *GAL4VP16* was used due to its short length (Asakawa et al., 2008). We used sgRNA sequences of GAAAAAGGGAAAGTTACTGGG for targeting the *tac1* gene to establish *Tg(tac1:zGFF)*, and of TGTCCATCTGCAGCTTATGGG for targeting the *penkb* gene to establish *Tg(penkb:zGFF)*, so that the coding sequence of the zGFF and poly-A tail was inserted at the start codon of the tac1 and penkb genes. The inserted sequence in the *Tg(tac1:zGFF)* also contained *zpc0.5:GFP* (Onichtchouk et al., 2003) after the poly-A tail to label transgene-positive oocytes and embryos for ease of fluorescent screening. The *Tg(pitx2-hs:loxP-mCherry-loxP-zGFF)*, *Tg(crhb-hs-loxP-RFP-loxP-GFP)*, and *Tg(crhb-hs:GFP)* were established through the knock-in strategy which was previously published (Kimura et al., 2014). The *Tg(crhb-hs:GFP)* line was generated by crossing the *Tg(crhb-hs:loxP-RFP-loxP-GFP)* line with a ubiquitous-Cre fish.

#### *In situ* hybridization

*In situ* hybridization was performed on 120 µm vibratome sections of the adult brain and observed as described previously (Amo et al., 2010). *Solute carrier family 17 member 7a (slc17a7a/vglut1), solute carrier family 17 member 6b (slc17a6b/vglut2.1), solute carrier family 17 member 6a (slc17a6a/vglut2.2), glutamate decarboxylase 1b (gad1b/gad67), glutamate decarboxylase 2 (gad2/gad65), tachykinin precursor 1 (tac1), proenkephalin b (penkb), NK2 homeobox 1 (nkx2.1), Neuropeitide Y (npy), prepronociceptin a (pnoca), neurotensin (nts), corticotropin releasing hormone b (crhb)* and *special AT-rich sequence binding protein homeobox 1b (satb1b)* were used as probes. Two-color fluorescent *in situ* hybridizations were performed on 70 µm vibratome sections of the adult brain and observed as described previously (Amo et al., 2010). A digoxigenin-labeled probe for *crhb* and a fluorescein-labeled probe for *npy* were visualized with the TSA Plus System (PerkinElmer) and FastRed (Roche), respectively.

#### Immunohistochemistry

Immunohistochemistry was performed on 75 µm vibratome coronal slices of the adult brain. All the primary and secondary antibodies used in this study are listed in the Key Resource Table. Some slices were counterstained with DAPI (Invitrogen) or fluorescent Nissl staining by Neurotrace-640/660 or Neurotrace-500/525 (Invitrogen). For permeabilization, phosphate-buffered saline (PBS) + 0.1% Triton-X was used throughout the primary and secondary antibody reactions in the regular condition. When immunohistochemistry was combined with DiI tracing, PBS + 0.3% Tween-20 was used only for 15 min before the primary antibody treatment so as not to dissolve the DiI staining in the buffer. During the primary antibody treatment, 1% of blocking reagent (Roche) was also added to the buffer.

The obtained slices were observed by commercial confocal laser scanning microscopy (Nikon C2+ or Zeiss LSM510) with 5 µm or 2.5 µm intervals of each optical section. Maximum intensity projection of all the optical sections from a slice (75 µm thick) was used for lower magnification images to show the entire view of the slice. A single optical section was shown for higher magnification images to show co-localization or segregation of fluorescent signals.

#### DiI tracing

A crystal of DiI (1,1’-Dioctadecyl-3,3,3’,3’-Tetramethylindocarbocyanine Perchlorate, Invitrogen) or CellTracker CM-DiI Dye (Invitrogen) is inserted with forceps into a target brain region which is exposed to a coronal cross section by a vibratome. Then the brain sample was soaked with PBS with 4% PFA and incubated at 37 degrees Celsius for 5 to 7 days. As for the experiment of Figure 4E-G, the incubation was only for 2 days to avoid too much spreading of diffused DiI from the application site. After incubation, the brain samples were sliced by the vibratome and underwent immunohistochemistry as described in the immunohistochemistry section.

#### Virus construction and preparation

For anterograde tracing, replication-competent vesicular stomatitis virus (VSV)-mCherry (1.0×1012 IU/ml) was used. The VSV vector, pVSV-mCherry, was created by using pVSV Venus VSVG (Addgene plasmid # 36399) (Beier et al., 2011) with the replacement of the first Venus position to mCherry. The VSV support plasmids pCAG-VSV-N, -P, and -L were gifts from Ian Wickersham (Addgene plasmid # 64087, # 64088, and # 64085). Virus supernatants were collected at 24-hour intervals and ultracentrifuged. The virus titer was determined by performing a dilution rate (Wickersham et al., 2010). The HSV vector, LT-HSV-EGFP (RN800; 5.0 x 10^9^ iu/ml), was used for retrograde tracing, obtained from the Massachusetts General Hospital (MGH) Gene Delivery Technology Core in the USA.

#### Virus injection

Wild-type or transgenic fish were continuously perfused and anesthetized with 0.02% tricaine (ethyl 3-aminobenzoate methanesulfonate salt, Sigma-Aldrich), and fish were mounted in a hand-made holder. A small opening in the skull was made with a micro drill above the coordinates for target areas (omniDrll35, WPI). 50 to 100 nl virus solution was injected at 125 um depth in the Dc and 225 um depth in the tectum using a micropump (UltraMicroPump II and Micro4, WPI). Then the orifice was covered with an ionized Teflon sheet, and the fish was allowed to recover with perfusion of teleost Ringer’s solution (Aoki et al., 2013). For tracing experiments by VSV, the fish was kept in a small chamber for 3-5 days at room temperature and then sacrificed. Especially in HSV-mediated tracing experiments, the fish was kept for seven days at 35℃.

#### Dissection, cell dissociation, and cell sorting for scRNAseq analysis

Dissection was performed on 6 individuals of *TgBAC(npy:GAL4VP16);Tg(UAS:GFP)* fish and 12 dissected tissue pieces from the left and right hemispheres were obtained in total. The dEN and its surrounding regions were carefully dissected with micro-scissors and fine forceps in ice-cold and oxygenized Neurobasal medium (ThermoFisher Scientific 21103049) supplemented with 1x B-27 (ThermoFisher Scientific 17504044) under a fluorescent dissection microscope, as shown in the Figure S5A. The dissected tissue was dissociated with the Papain Dissociation Kit (Worthington; LK003150) with 0.1% 2-mercaptoethanol for 15 minutes with gentle shaking at 28.5 degrees Celsius. Then, the cells were dissociated by gentle trituration 15 times with a glass Pasteur pipet coated with 2% BSA in PBS and spun at 300xg for 5 minutes. The cells were resuspended in papain inhibitor solution (Worthington) and incubated for 10 minutes with gentle shaking at 28.5 degrees Celsius. Then, the cells were further dissociated by gentle trituration 20 times with a glass Pasteur pipet attached with a regular 200-µl tip coated with 2% BSA in PBS. The dissociated cell suspension was then filtered with pluriStrainer Mini 40 µm (pluriSelect) coated with 2% BSA in PBS and spun at 300xg for 5 minutes. The resulting cell suspension was resuspended in 2% BSA in PBS, and then cell debris and dead cells were removed by FACS (FACSAria SORP, BD Biosciences) using Hoechst (to sort out cells from cell debris) and Propidium Iodide (to sort out living cells from dead cells). After FACS sorting, a small fraction of the cell suspension was used to estimate the total number of the cells and their viability using a dead cell stain Trypan Blue (elabscience). The obtained suspension contained 16,000 cells with 85.0% viability.

The resulting single-cell suspension was loaded on the Chromium Next GEM Single Cell 3’ Reagent Kits v3.1 (10x Genomics, PN-1000269), and the cDNA library was prepared according to the manufacturer’s instructions. The obtained cDNA library underwent Next generation sequencing by illumina Hiseq X (GENEWIZ) with 400,429,716 total reads and 85.9% of sequencing saturation. The obtained sequence was then analyzed by “Cell Ranger count” pipeline provided by 10x Genomics with default options. The reads were aligned to zebrafish reference transcriptome (ENSEMBL Zv11, release 99) and EGFP CDS, which were built by “Cell Ranger mkref” command based on zebrafish reference genome GRCz11 and annotation Ensembl 99. This resulted in 3381 estimated number of cells with 4890 median unique molecular identifier (UMI) counts per cell.

#### Data Analysis by Seurat

The obtained scRNAseq data of 3381 cells were analyzed by Seurat (Butler et al., 2018). As a quality control, cells with more than 6% mitochondrial genes, less than 200 unique genes, and more than 17500 UMIs were removed. Barcodes with less than 500 UMIs had been already removed by cellranger count pipeline. The remaining 3043 cells were then data normalized by LogNormalize method with scale.factor = 10000.

For further calculation of UMAP and clustering, variable genes were determined with FindVariableFeatures function with selection.method = “vst”, and the top 2000 most highly variable genes were used for further clustering. The expression of each gene was shifted so that the mean expression across cells is 0, and scaled so that the variance across cells is 1 by ScaleData function. PCA was run on the scaled data and then UMAP and clustering were performed with the FindNeighbors function with the top 35 PCs, the FindClusters function with resolution = 0.5, and the RunUMAP function with the top 35 PCs. Dot plots and violin plots were generated by Seurat and cell types were determined by the expression of marker genes that define specific cell types.

#### Supplemental information

Document S1. Figures S1-S8 and Tables S1-3.

Data S1. List of differentially expressed genes in the individual cell clusters, related to Figure 6.

Data S2. List of differentially expressed genes in the [*npy-*negative and *gad1b*-positive] cells and [*npy*-positive] cells in the EN clusters, related to Figure 6.

