## Supplementary material for "Transgenic tools targeting striatal and pallidal subpopulations revealed evolutionary conservation and specialization of the cortico-basal ganglia circuit in zebrafish": Tanimoto_et_al_Supplementary_Information

Figure S1

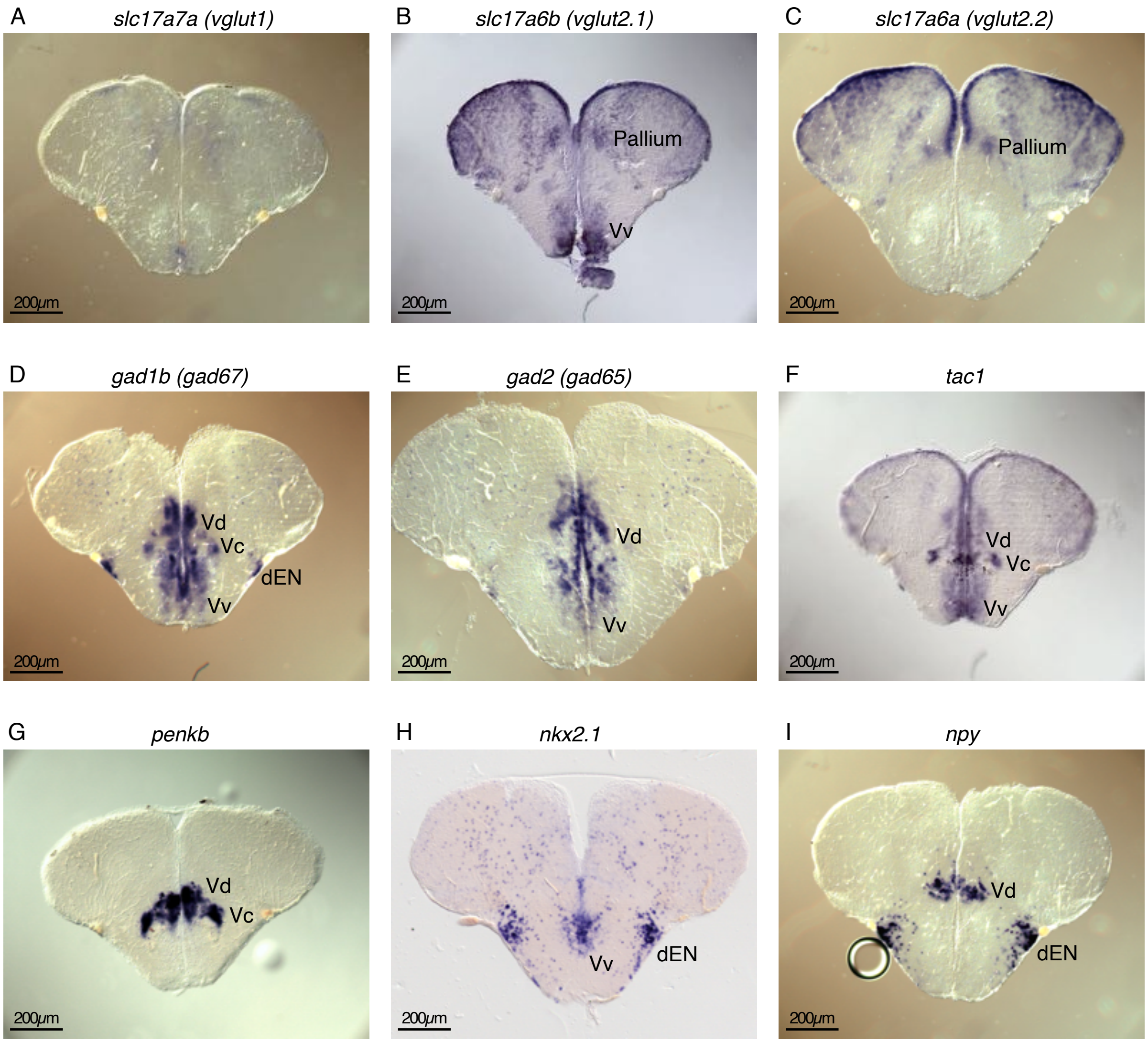

A
*slc17a7a (vglut1)*
200μm
B
*slc17a6b (vglut2.1)*
Pallium
Vv
200μm
C
*slc17a6a (vglut2.2)*
Pallium
200μm
D
*gad1b (gad67)*
Vd
Vc
dEN
Vv
200μm
E
*gad2 (gad65)*
Vd
Vv
200μm
F
*tac1*
Vd
Vc
Vv
200μm
G
*penkb*
Vd
Vc
200μm
H
*nkx2.1*
Vv
dEN
200μm
I
*npy*
Vd
dEN
200μm

**Figure S1. Gene expression profiles of the pallium and the subpallial regions (Related to Figure 1)**

(A-I) *In situ* hybridization analysis of glutamatergic neuronal markers (*slc17a7a* (*vglut1*), *slc17a6b* (*vglut2.1*), and *slc17a6a* (*vglut2.2*)), GABAergic neuronal markers (*gad1b* (*gad67*) and *gad2* (*gad65*)), striatal neuronal markers (*tac1* and *penkb*), and pallidal neuronal markers (*nkx2.1* and *npy*). Coronal sections in the anterior telencephalon are shown.

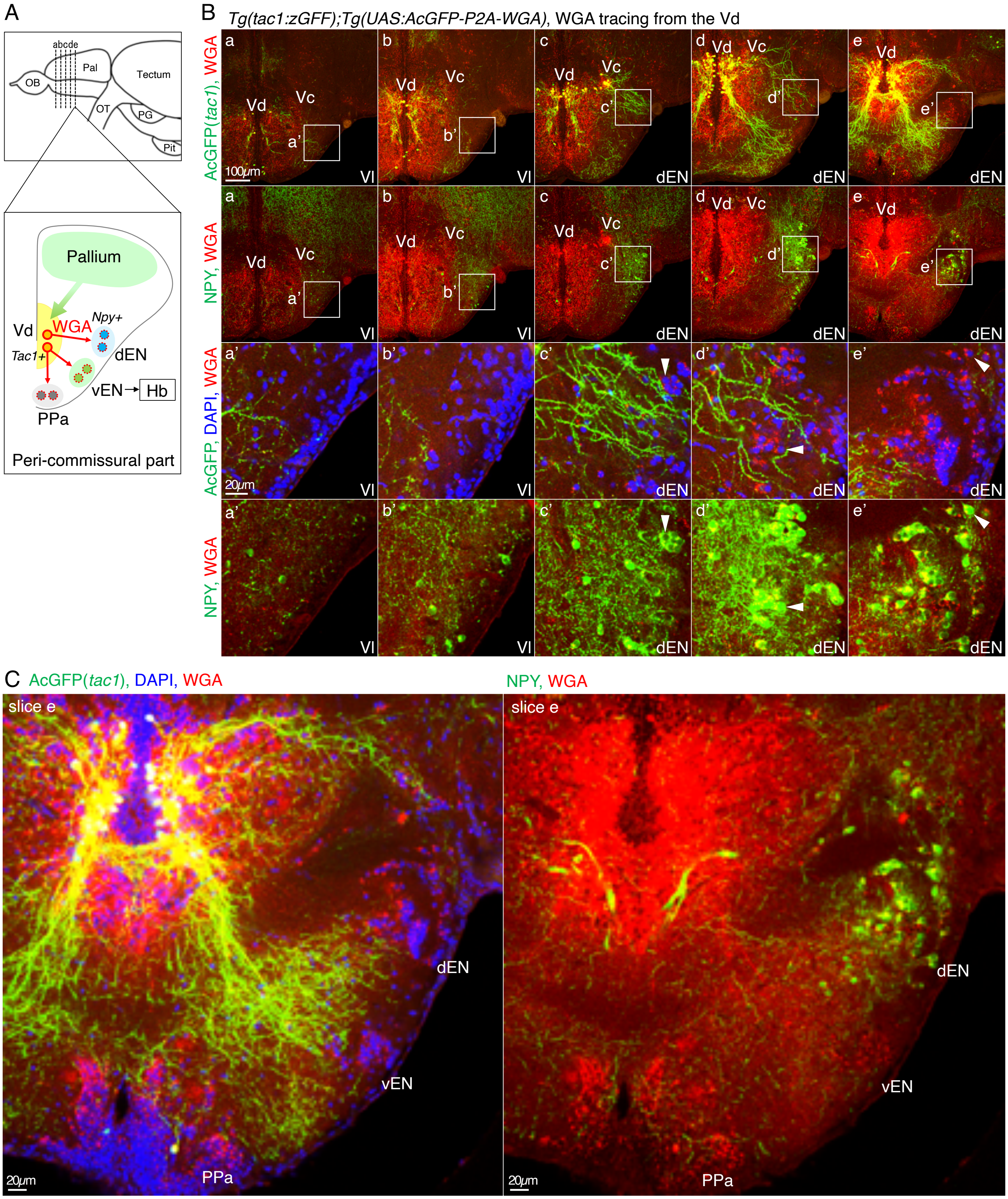
A
abcde
OB
Pal
Tectum
OT
PG
Pit
Pallium
Vd
WGA
Npy+
Tac1+
dEN
vEN
Hb
PPa
Peri-commissural part
B
Tg(tac1:zGFF);Tg(UAS:AcGFP-P2A-WGA), WGA tracing from the Vd
AcGFP(tac1), WGA
NPY, WGA
AcGFP, DAPI, WGA
NPY, WGA
a
b
c
d
e
Vd
Vc
a'
b'
c'
d'
e'
Vl
dEN
100μm
20μm
C
AcGFP(tac1), DAPI, WGA
NPY, WGA
slice e
dEN
vEN
PPa
20μm

**Figure S2. WGA tracing from the *tac1*+ Vd striatal neurons caused WGA accumulation in the dEN, vEN, and PPa (Related to Figure 2)**

(A) Illustration of a coronal slice depicting WGA tracing from the *tac1*+ Vd neurons and trans-synaptic WGA transfer to the dEN, vEN, and PPa at the peri-commissural telencephalon. The top panel illustrates the lateral view of the zebrafish brain, and the dashed lines indicate antero-posterior positions of the coronal slices (a-e) shown in the panel B.

(B) WGA expression in the *tac1*+ Vd neurons and immunohistochemistry of AcGFP (green), WGA (red), DAPI (blue), and NPY (green). Five successive coronal slices (a-e) are shown so that the first two slices contain the NPY-negative Vl region and the latter three slices contain the NPY+ dEN region. Insets in the panels a-e (top two rows) show the positions of the panels a'-e' (bottom two rows), focusing on the Vl or dEN. In the panels c'-e', arrowheads indicate representative NPY+ dEN neurons with WGA signals.

(C) Magnified overviews of the slice e in B. Innervation of AcGFP+ fibers and WGA accumulation occurred in the dEN, vEN, and PPa.

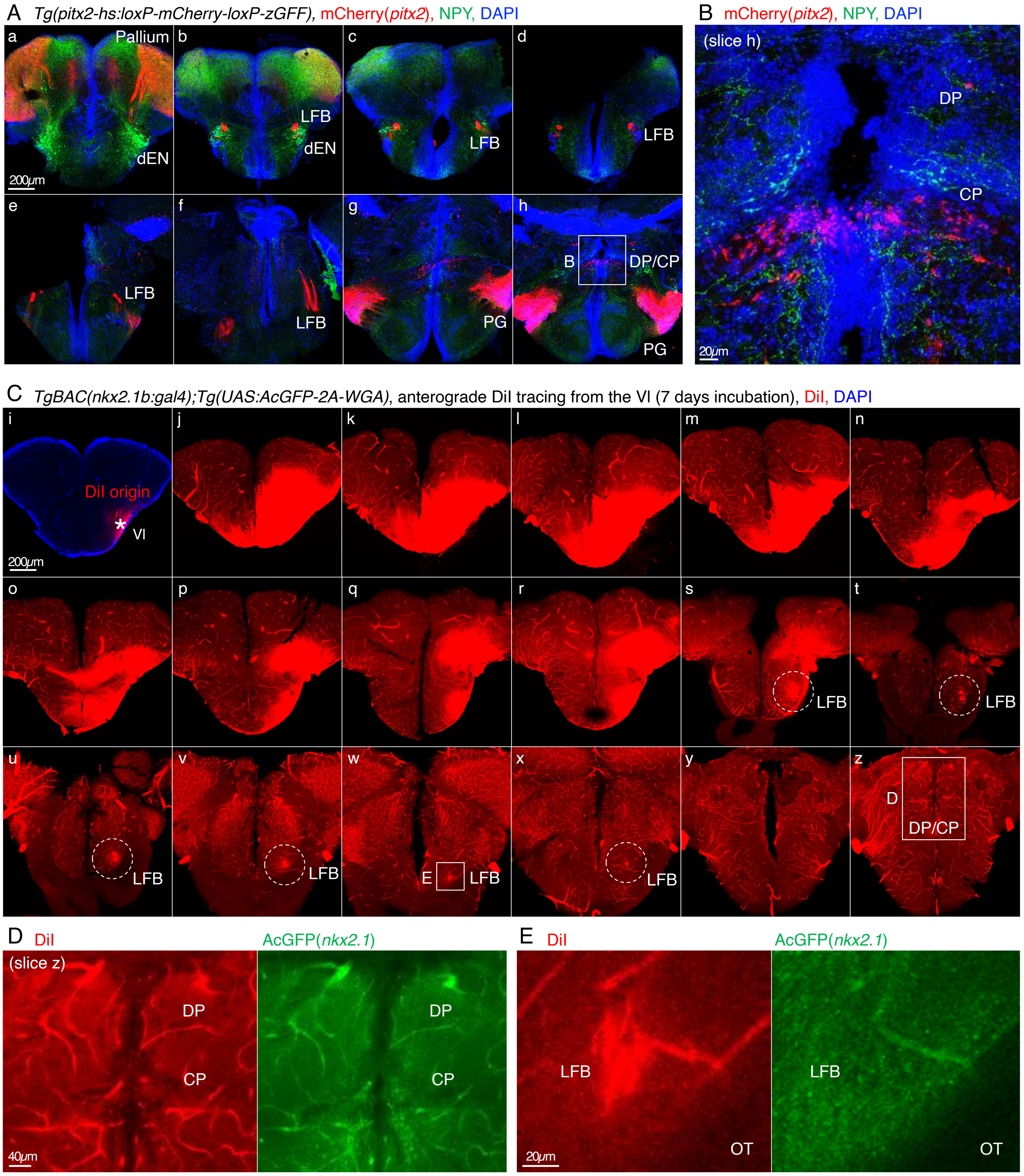
A Tg(pitx2-hs:loxP-mCherry-loxP-zGFF), mCherry(pitx2), NPY, DAPI
a Pallium dEN 200μm
b LFB dEN
c LFB
d LFB
e LFB
f LFB
g PG
h B DP/CP PG
B mCherry(pitx2), NPY, DAPI
(slice h)
DP
CP
20μm
C TgBAC(nkx2.1b:gal4);Tg(UAS:AcGFP-2A-WGA), anterograde DiI tracing from the Vl (7 days incubation), DiI, DAPI
i DiI origin Vl 200μm
j
k
l
m
n
o
p
q
r
s LFB
t LFB
u LFB
v LFB
w E LFB
x LFB
y
z D DP/CP
D DiI AcGFP(nkx2.1)
(slice z)
DP
CP
40μm
E DiI AcGFP(nkx2.1)
LFB
OT
20μm

**Figure S3. The indirect-pathway targeted region Vl seemed not to project to the possible teleost subthalamic regions (Related to Figure 3)**

(A) Expression pattern analysis of *Tg(pitx2-hs:loxP-mCherry-loxP-zGFF)*. Immunohistochemistry of mCherry (red), NPY (green), and DAPI (blue). Eight successive coronal slices (a-h) from the peri-commissural telencephalon to the diencephalon are shown. An inset in the panel h shows the position of the panel B, focusing on the expression in the DP/CP region. Note that the PG also contained *pitx2*+ neurons, which projected to the pallium via the LFB.

(B) A magnified view of the DP/CP in the slice h. A part of the CP showed expression of mCherry by the promoter of the mammalian STN marker *pitx2*.

(C) DiI tracing from the Vl anterogradely labeled projection fibers in the ipsilateral LFB. *TgBAC(nkx2.1b:gal4);Tg(UAS:AcGFP-2A-WGA)* fish was used to apply DiI to the Vl correctly. Eighteen successive coronal slices (i-z) from the Vl region to the diencephalon are shown. An asterisk in the panel i indicates DiI origin. Dotted circles in the panels s-x indicate DiI+ projection fibers in the LFB. Insets in the panel z and w show the positions of the panels D and E, respectively.

(D) Magnified views of the DP/CP in the slice z. Anterograde DiI tracing from the Vl did not clearly stain any projection fibers around the CP.

(E) Magnified views of the LFB in the slice w. Immunohistochemistry of AcGFP (green) is combined with DiI tracing (red). The DiI-stained fibers which originates from the Vl and exit from the telencephalon are not co-stained with AcGFP expressed by the *Nkx2.1* promoter, suggesting that the *Nkx2.1*+ neurons in the Vl mostly project intra-telencephalically. OT, optic tract.

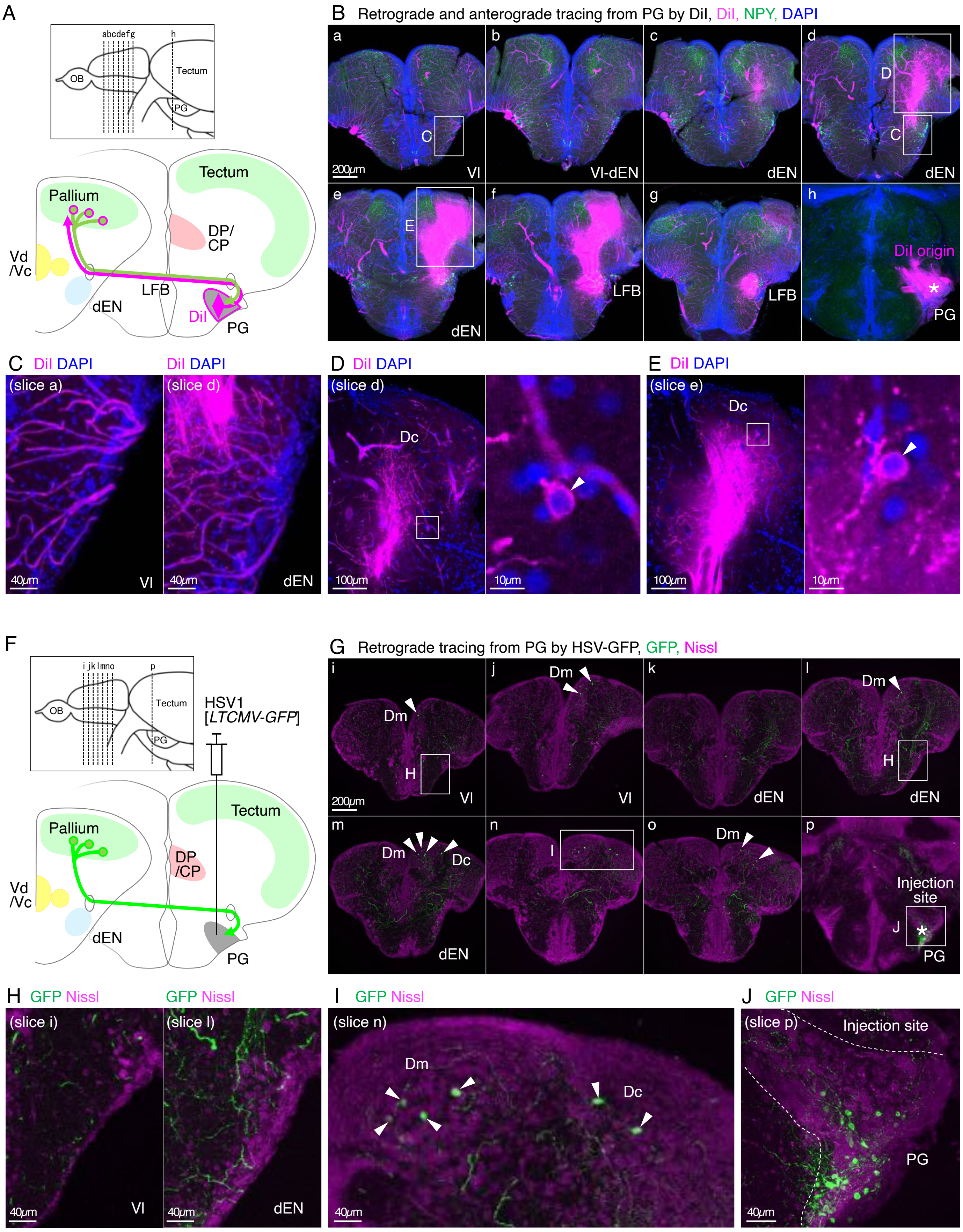

A
abcdefg
h
OB
Tectum
PG
Tectum
Pallium
DP/
CP
Vd
/Vc
LFB
dEN
DiI
PG
B Retrograde and anterograde tracing from PG by DiI, DiI, NPY, DAPI
a
b
c
d
C
200μm
Vl
Vl-dEN
dEN
D
C
dEN
e
f
g
h
E
DiI origin
*
PG
dEN
LFB
LFB
C DiI DAPI
(slice a)
40μm
Vl
DiI DAPI
(slice d)
40μm
dEN
D DiI DAPI
(slice d)
Dc
100μm
10μm
E DiI DAPI
(slice e)
Dc
100μm
10μm
F
i jklmno
p
OB
Tectum
PG
HSV1
[LTCMV-GFP]
Tectum
Pallium
DP
/CP
Vd
/Vc
dEN
PG
G Retrograde tracing from PG by HSV-GFP, GFP, Nissl
i
Dm
H
200μm
Vl
j
Dm
Vl
k
dEN
l
Dm
H
dEN
m
Dm
Dc
dEN
n
I
o
Dm
p
Injection
site
J
*
PG
H GFP Nissl
(slice i)
40μm
Vl
GFP Nissl
(slice l)
40μm
dEN
I GFP Nissl
(slice n)
Dm
Dc
40μm
J GFP Nissl
(slice p)
Injection site
PG
40μm

**Figure S4. Retrograde DiI or HSV virus tracing from the PG did not label any neurons in the dEN (Related to Figure 5)**

(A-B) DiI tracing from the PG. Immunohistochemistry of NPY (green) is combined with DiI tracing (magenta) to mark the dEN. Seven successive coronal slices at the telencephalon (a-g) and one coronal slice at the PG (h) are shown. Insets in the panels a, d, and e show the positions of the panels C-E. An asterisk in the panel h indicates DiI origin.

(C) Magnified views of the Vl in the slice a (left panel) and the dEN in the slice d (right panel). DiI tracing from the PG did not retrogradely label any neurons in the dEN, and therefore the PG seemed not to correspond to the motor thalamus.

(D) Magnified views of the pallium in the slice d. DiI tracing from the PG anterogradely labeled projection fibers in the pallium (left panel), and retrogradely labeled pallial neurons (right panel). An inset in the left panel is magnified in the right panel. An arrowhead in the right panel indicates a retrogradely labeled pallial neuron.

(E) Same as D, but in the slice e.

(F-G) HSV-GFP injection to the PG and immunohistochemistry of GFP (green) and Nissl (magenta). Seven successive coronal slices at the telencephalon (i-o) and one coronal slice at the PG (p) are shown. Arrowheads indicate retrogradely labeled pallial neurons. Insets in the panels i, l, n, and p show the positions of the panels H-J. An asterisk in the panel p indicates an approximate injection point.

(H) Magnified views of the Vl in the slice i (left panel) and the dEN in the slice l (right panel). HSV-GFP injection to the PG did not retrogradely label any neurons in the dEN, and therefore the PG seemed not to correspond to the motor thalamus.

(I) A magnified view of the pallium in the slice n. HSV-GFP injection to the PG retrogradely labeled pallial neurons (arrowheads).

(J) A magnified view of the injection site of the PG in the slice p. Dotted line indicates an outline of the PG region.

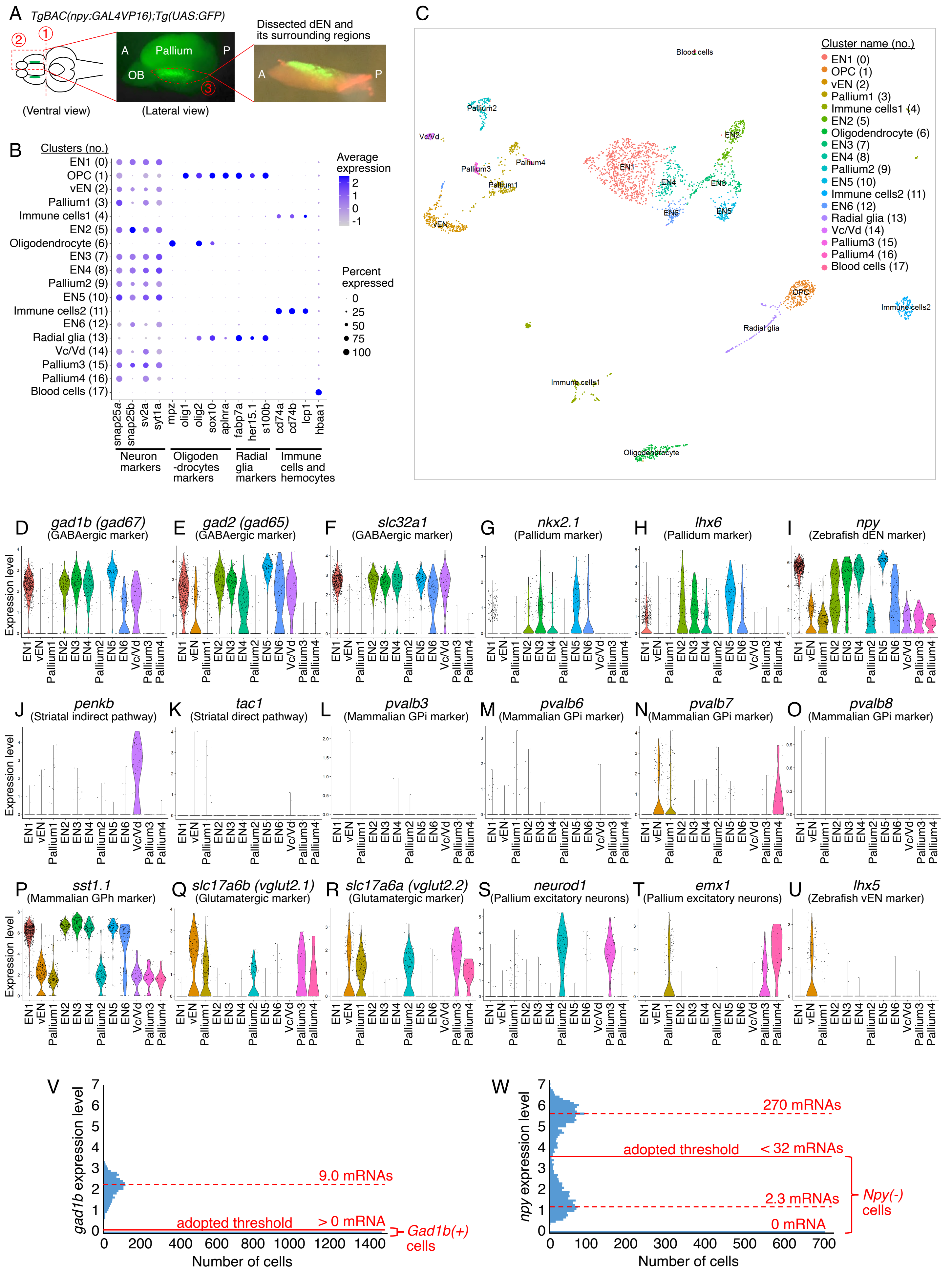
A TgBAC(npy:GAL4VP16);Tg(UAS:GFP)
(Ventral view)
(Lateral view)
Dissected dEN and its surrounding regions
Pallium
OB
B
Clusters (no.)
EN1 (0)
OPC (1)
vEN (2)
Pallium1 (3)
Immune cells1 (4)
EN2 (5)
Oligodendrocyte (6)
EN3 (7)
EN4 (8)
Pallium2 (9)
EN5 (10)
Immune cells2 (11)
EN6 (12)
Radial glia (13)
Vc/Vd (14)
Pallium3 (15)
Pallium4 (16)
Blood cells (17)
Average expression
Percent expressed
Neuron markers
Oligoden-drocytes markers
Radial glia markers
Immune cells and hemocytes
C
Cluster name (no.)
D gad1b (gad67) (GABAergic marker)
E gad2 (gad65) (GABAergic marker)
F slc32a1 (GABAergic marker)
G nkx2.1 (Pallidum marker)
H lhx6 (Pallidum marker)
I npy (Zebrafish dEN marker)
J penkb (Striatal indirect pathway)
K tac1 (Striatal direct pathway)
L pvalb3 (Mammalian GPi marker)
M pvalb6 (Mammalian GPi marker)
N pvalb7 (Mammalian GPi marker)
O pvalb8 (Mammalian GPi marker)
P sst1.1 (Mammalian GPh marker)
Q slc17a6b (vglut2.1) (Glutamatergic marker)
R slc17a6a (vglut2.2) (Glutamatergic marker)
S neurod1 (Pallium excitatory neurons)
T emx1 (Pallium excitatory neurons)
U lhx5 (Zebrafish vEN marker)
Expression level
V
gad1b expression level
9.0 mRNAs
adopted threshold
> 0 mRNA
Gad1b(+) cells
Number of cells
W
npy expression level
270 mRNAs
adopted threshold
< 32 mRNAs
2.3 mRNAs
0 mRNA
Npy(-) cells
Number of cells

**Figure S5. ScRNAseq analysis of the zebrafish pallidum and characterization of each cell cluster obtained from the unbiased clustering analysis (Related to Figure 6)**

(A) Dissection of the dEN and its surrounding regions from the brain of *TgBAC(npy:GAL4VP16);Tg(UAS:GFP)* fish. Red dotted lines are carefully cut by microscissors and forceps in the designated order.

(B) Dot plots showing expression patterns of neuron- and various non-neuron-markers in all the detected clusters. Dot color indicates average expression levels and dot size indicates percent expressed. Based on this result, clusters no. 1, 4, 6, 11, 13, and 17 are determined as non-neuronal clusters of oligodendrocytes precursor cells (OPC), immune cells1, oligodendrocytes, immune cells2, radial glia, and blood cells, respectively.

(C) UMAP plot of all the clusters including non-neuronal ones.

(D-U) Violin plots showing natural log-normalized expression levels of various neuronal markers in the 12 neuronal clusters. *Pvalb1, pvalb2, pvalb4, pvalb5*, and *pvalb9* were also tested, but not expressed at all.

(V-W) Histograms of expression levels of *gad1b* (panel V) and *npy* (panel W). Distribution of *gad1b* expression levels showed a single peak at 9.0 mRNAs and accumulation of data point at 0 mRNA, and thus >0 mRNA was chosen as a threshold to determine *gad1b*-positive cells. Distribution of *npy* expression levels showed two peaks at 270 mRNAs and at 2.3 mRNAs, and thus the minima between the two peaks (<32 mRNAs) was chosen as a threshold to determine *npy*-negative cells. The lower peak at 2.3 mRNAs may have been caused by background contamination of *npy* mRNAs released from dead cells because of its extraordinarily high expression levels up to ~1000 mRNAs. The numbers of mRNAs are per 10000 mRNAs.

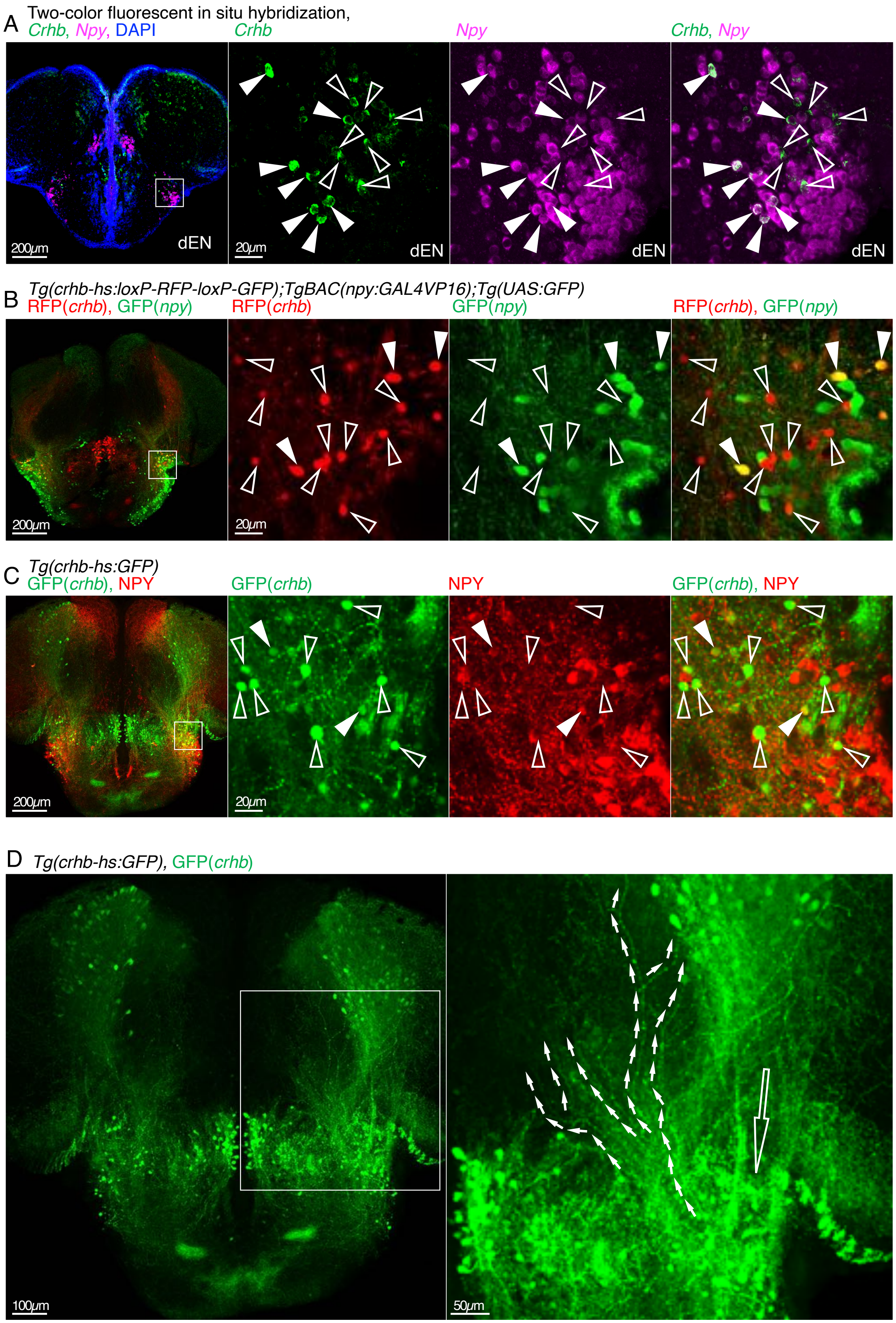
A
Two-color fluorescent in situ hybridization,
Crhb, Npy, DAPI
Crhb
Npy
Crhb, Npy
200μm
20μm
dEN
B
Tg(crhb-hs:loxP-RFP-loxP-GFP);TgBAC(npy:GAL4VP16);Tg(UAS:GFP)
RFP(crhb), GFP(npy)
RFP(crhb)
GFP(npy)
RFP(crhb), GFP(npy)
200μm
20μm
C
Tg(crhb-hs:GFP)
GFP(crhb), NPY
GFP(crhb)
NPY
GFP(crhb), NPY
200μm
20μm
D
Tg(crhb-hs:GFP), GFP(crhb)
100μm
50μm

**Figure S6. Expression pattern analysis of the *crhb* and the transgenic labeling by the *crhb* promoter (Related to Figure 6)**

(A) Two-color fluorescent in situ hybridization of *crhb* (green), *npy* (magenta), and DAPI (blue). An inset in the leftmost panel is magnified in the right three panels. In the right three panels, filled arrowheads indicate *npy* and *crhb* double-positive neurons, and open arrowheads indicate *crhb* only-positive neurons.

(B) Immunohistochemistry of *crhb*+ neurons (RFP, red) and *npy*+ neurons (GFP, green) in *Tg(crhb-hs:loxP-RFP-loxP-GFP);TgBAC(npy:GAL4VP16);Tg(UAS:GFP)* fish. An inset in the leftmost panel is magnified in the right three panels. Again, filled arrowheads indicate *npy* and *crhb* double-positive neurons, and open arrowheads indicate *crhb* only-positive neurons.

(C) Immunohistochemistry of *crhb*+ neurons (GFP, green) and NPY+ neurons (NPY, red) in *Tg(crhb-hs:GFP)* fish. An inset in the leftmost panel is magnified in the right three panels. Again, filled arrowheads indicate *npy* and *crhb* double-positive neurons, and open arrowheads indicate *crhb*-only positive neurons.

(D) Detailed expression patterns in the telencephalon of *Tg(crhb-hs:GFP)* fish. Immunohistochemistry of *crhb*+ neurons and fibers (GFP, green). The same animal as panel C. An inset in the left panel is magnified in the right panel. In the right panel, a large open arrow indicates *crhb*+ projection bundles, which putatively originate from the pallial *crhb*+ neurons. Small filled arrows indicate *crhb*+ projection fibers, which putatively originate from the *crhb*+ dEN neurons and extend toward the pallial regions.

Figure S7

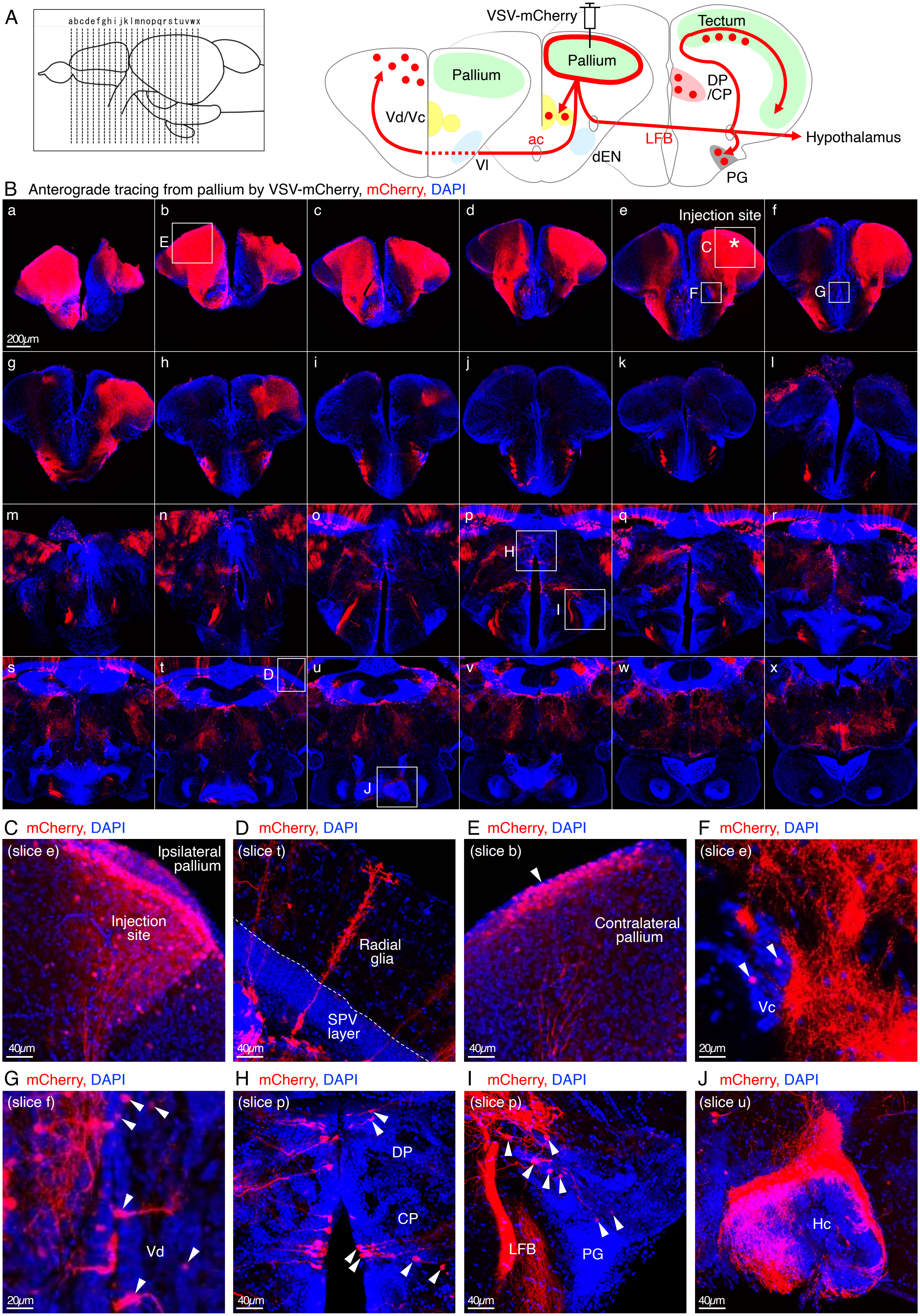

A
abcdefghijklmnopqrstuvwx
VSV-mCherry
Pallium
Pallium
Tectum
Vd/Vc
DP
/CP
ac
LFB
Hypothalamus
Vl
dEN
PG
B Anterograde tracing from pallium by VSV-mCherry, mCherry, DAPI
a
b
c
d
e
f
E
Injection site
C
F
G
200μm
g
h
i
j
k
l
m
n
o
p
q
r
H
I
s
t
u
v
w
x
D
J
C mCherry, DAPI
(slice e)
Ipsilateral
pallium
Injection
site
40μm
D mCherry, DAPI
(slice t)
Radial
glia
SPV
layer
40μm
E mCherry, DAPI
(slice b)
Contralateral
pallium
40μm
F mCherry, DAPI
(slice e)
Vc
20μm
G mCherry, DAPI
(slice f)
Vd
20μm
H mCherry, DAPI
(slice p)
DP
CP
40μm
I mCherry, DAPI
(slice p)
LFB
PG
40μm
J mCherry, DAPI
(slice u)
Hc
40μm

**Figure S7. Visualization of output pathways from the pallium to the other parts of the brain (Related to Figure 7)**

(A-B) Same as Figure 7A-B, but with extended successive coronal slices (k-o and q-x). Immunohistochemistry of mCherry (red), and DAPI (blue). An asterisk in the panel e indicates an approximate injection point. Insets show the positions of the panels C-J.

(C) A magnified view of the injection site in the ipsilateral pallium in the slice e.

(D) A magnified view of single radial glia showing the "bottlebrush-like" morphology. Dotted line indicates the boundary between the SPV and the upper layer.

(E-J) Magnified views of the contralateral pallium, Vc, Vd, DP/CP, PG, and Hc. Arrowheads indicate representative anterogradely labeled mCherry+ pallial neurons.

Figure S8

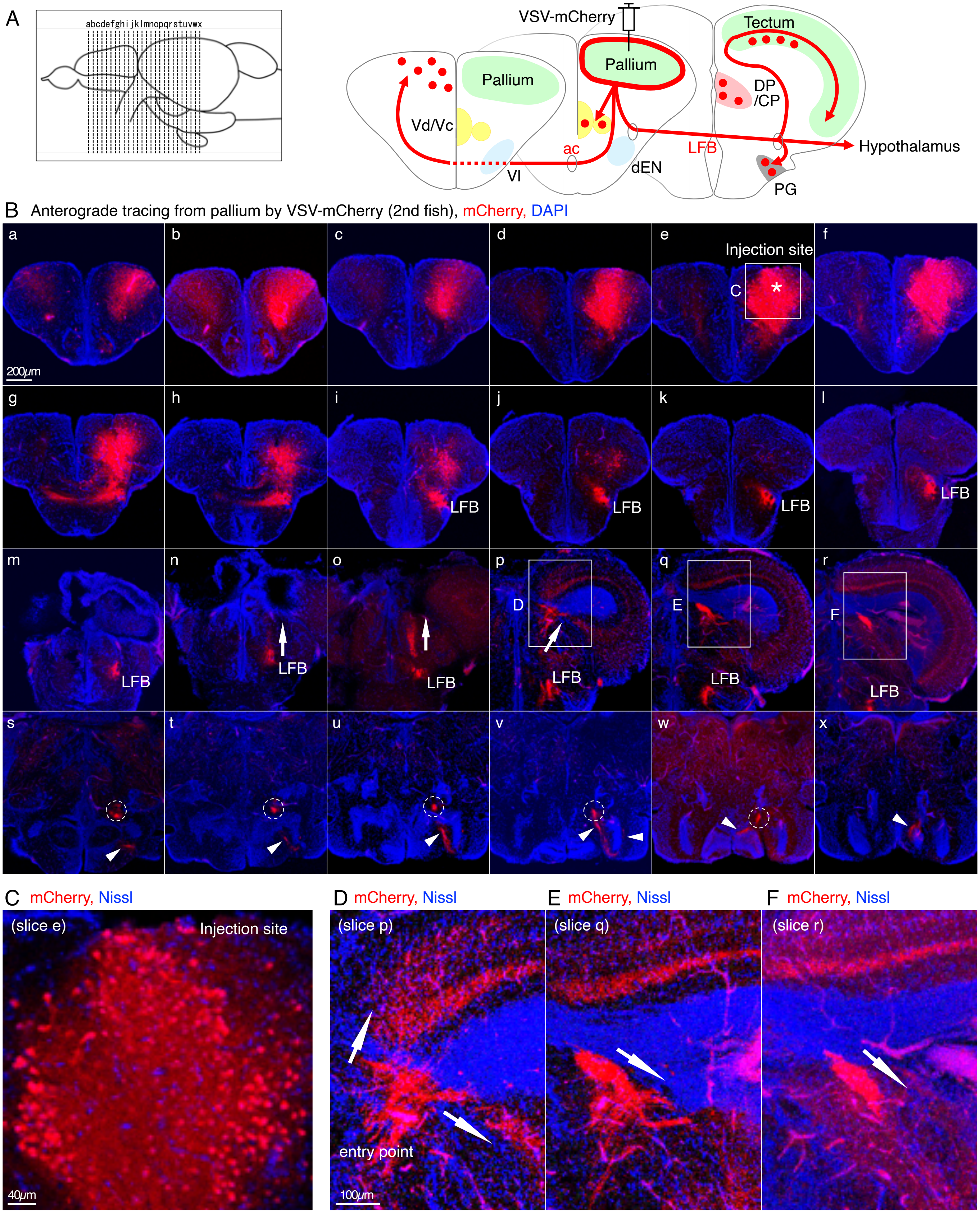
A
abcdefghijklmnopqrstuvwx
VSV-mCherry
Tectum
Pallium
Pallium
DP
/CP
Vd/Vc
ac
LFB
Hypothalamus
Vl
dEN
PG
B Anterograde tracing from pallium by VSV-mCherry (2nd fish), mCherry, DAPI
a
b
c
d
e
f
Injection site
C
200μm
g
h
i
j
k
l
LFB
m
n
o
p
q
r
D
E
F
s
t
u
v
w
x
C mCherry, Nissl
(slice e)
Injection site
40μm
D mCherry, Nissl
(slice p)
entry point
100μm
E mCherry, Nissl
(slice q)
F mCherry, Nissl
(slice r)

**Figure S8. Same as Figure S7, but in the 2nd example with a lower level of trans-synaptic virus infection (Related to Figure 7)**

(A-B) VSV-mCherry injection into the pallium of another fish visualized entering pallio-tectal projection fibers from the LFB to the ipsilateral tectum. Immunohistochemistry of mCherry (red) and Nissl (blue). Twenty-four successive coronal slices were shown. An asterisk in the panel e indicates an approximate injection point. In this fish, most of the mCherry+ neurons were located around the injection site in the ipsilateral pallium (panels a-j). The mCherry+ projection fibers from these pallial neurons extended to the extra-telencephalic area solely through the ipsilateral LFB (panels k-m). Then, the mCherry+ projection fiber bundles were bifurcated into two branches, one remained in the LFB and the other climbed up toward the tectum (panels n-p, arrows). The remained projection fibers in the LFB extended to the hypothalamic regions including the Hc (panels s-x, dotted circles and arrowheads). Insets in the panels e and p-r show the positions of C and D-F, respectively.

(C) A magnified view of the injection site in the ipsilateral pallium in the slice e.

(D-F) Magnified views of the entry point of the pallio-tectal projection fibers in the slices p-r. Arrows indicate the directions of the entering fibers to the presynaptic layers of the tectum.

### Supplementary tables

| Targeted gene | *Tachykinin Precursor 1* | | *Proenkephalin b* | *NK2 homeobox 1* | *Neuropeptide Y* | *Glutamate decarboxylase 1b* |
|---|---|---|---|---|---|---|
| Gal4 driver line | *TgBAC(tac1:GAL4VP16)*$^{rw157}$ | *Tg(tac1:zGFF)*$^{rw158}$ | *Tg(penkb:zGFF)*$^{rw159}$ | *TgBAC(nkx2.1:GAL4VP16)*$^{rw160}$ | *TgBAC(npy:GAL4VP16)*$^{rw161b}$ | *TgBAC(gad1b:GAL4VP16)*$^{rw162b}$ |
| Method | Transposon-mediated BAC | CRISPR-Cas9 knock-in | CRISPR-Cas9 knock-in | Transposon-mediated BAC | Transposon-mediated BAC | Transposon-mediated BAC |
| Main expression target(s) in the telencephalon | Vc, weakly in Vd, Vv, and a part of Dl | Vd | Vc, and weakly in Vd | Vl, dEN, Vv, and sparsely in Dl | dEN and partly in Vl | GABAergic neurons |
| Expression in the mammalian basal ganglia | Direct pathway striatal projection neurons | | Indirect-pathway striatal projection neurons | Prototypic GP/GPe neurons | Ventral pallidum | GABAergic neurons |

**Table S1. Transgenic tools to characterize anatomical properties of specific neuronal populations of the zebrafish basal ganglia (Related to Figure 1)**

| | Number of tested individuals | Total number of DiI+ dEN neurons | Total number of DiI+ anti-GFP+ dEN neurons | Percentage of anti-GFP+ neurons in all DiI+ neurons |
|---|---|---|---|---|
| *TgBAC(gad1b:GAL4VP16)* | 3 | 27 | 15 | 55.6 % |
| *TgBAC(npy:GAL4VP16)* | 3 | 7 | 0 | 0 % |
| *Tg(crhb-hs:GFP)* | 2 | 5 | 2 | 40.0 % |

**Table S2. Statistics of the retrogradely labeled cells in the dEN after DiI application to the DP/CP thalamus (Related to Figures 5 and 6)**

| | EN1 | EN2 | EN3 | EN4 | EN5 | EN6 |
|---|---|---|---|---|---|---|
| Number of [*npy*-negative and *gad1b*-positive] cells /total cells in each cluster | 58/771 cells (7.5 %) | 98/182 cells (53.8 %) | 47/166 cells (28.3 %) | 3/154 cells (1.9 %) | 1/124 cells (0.8 %) | 28/108 cells (25.9 %) |
| Number of [*npy*-negative and *gad1b*-positive] cells /total such cells in all the clusters | 58/235 cells (24.7 %) | 98/235 cells (41.7 %) | 47/235 cells (20.0 %) | 3/235 cells (1.3 %) | 1/235 cells (0.4 %) | 28/235 cells (11.9 %) |

**Table S3. Summary of the number of [*npy*-negative and *gad1b*-positive] cells in the six EN clusters obtained by scRNAseq analysis (Related to Figure 6)**
